# Two distinct ferredoxins are essential for nitrogen fixation by the iron nitrogenase in *Rhodobacter capsulatus*

**DOI:** 10.1101/2023.12.07.570605

**Authors:** Holly Addison, Timo Glatter, Georg K. A. Hochberg, Johannes G. Rebelein

**Author notes:** **Corresponding author:** Correspondence to Johannes G. Rebelein.

## Abstract

Nitrogenases are the only enzymes able to fix gaseous nitrogen into bioavailable ammonia and, hence, are essential for sustaining life. Catalysis by nitrogenases requires both a large amount of ATP and electrons donated by strongly reducing ferredoxins or flavodoxins. Our knowledge about the mechanisms of electron transfer to nitrogenase enzymes is limited: The electron transport to the iron (Fe)-nitrogenase has hardly been investigated. Here, we characterised the electron transfer pathway to the Fe-nitrogenase in *Rhodobacter capsulatus* via proteome analyses, genetic deletions, complementation studies and phylogenetics. Proteome analyses revealed an upregulation of four ferredoxins under nitrogen-fixing conditions reliant on the Fe-nitrogenase in a molybdenum nitrogenase knockout strain, compared to non-nitrogen-fixing conditions. Based on these findings, *R. capsulatus* strains with deletions of ferredoxin (*fdx*) and flavodoxin (*fld, nifF*) genes were constructed to investigate their roles in nitrogen fixation by the Fe-nitrogenase. *R. capsulatus* deletion strains were characterised by monitoring diazotrophic growth and Fe-nitrogenase activity *in vivo*. Only deletions of *fdxC* or *fdxN* resulted in slower growth and reduced Fe-nitrogenase activity, whereas the double-deletion of both *fdxC* and *fdxN* abolished diazotrophic growth. Differences in the proteomes of Δ*fdxC* and Δ*fdxN* strains, in conjunction with differing plasmid complementation behaviours of *fdxC* and *fdxN*, indicate that the two Fds likely possess different roles and functions. These findings will guide future engineering of the electron transport systems to nitrogenase enzymes, with the aim of increased electron flux and product formation.

**Importance:** Nitrogenases are essential for biological nitrogen fixation, converting atmospheric nitrogen gas to bioavailable ammonia. Production of ammonia by diazotrophic organisms, harbouring nitrogenases, is essential for sustaining plant growth. Hence, there is a large scientific interest in understanding the cellular mechanisms for nitrogen fixation via nitrogenases. Nitrogenases rely on highly reduced electrons to power catalysis, though we lack knowledge as to which proteins shuttle the electrons to nitrogenases within cells. Here, we characterised the electron transport to the iron (Fe)-nitrogenase in the model diazotroph *Rhodobacter capsulatus*, showing that two distinct ferredoxins are very important for nitrogen fixation despite having different redox centres. Additionally, our research expands upon the debate on whether ferredoxins have functional redundancy or perform distinct roles within cells. Here, we observe that both essential ferredoxins likely have distinct roles based on differential proteome shifts of deletion strains and different complementation behaviours.

## Introduction

Nitrogenases are the only known enzymes to perform nitrogen fixation, the biological reduction of molecular nitrogen (N_2_) to bioavailable ammonia (NH_3_). Nitrogenases are found in many microorganisms across the domains of bacteria to archaea^[1]^. Catalysis by nitrogenases requires chemical energy in the form of ATP and low-potential electrons to reduce N_2_. The optimal catalytic activity is shown in (1)^[2]^:

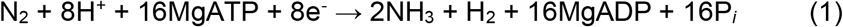

There are three known nitrogenase isoforms: the canonical molybdenum (Mo)-nitrogenase and two alternative nitrogenases, the vanadium (V)-nitrogenase and the iron (Fe)-nitrogenase (Fig. 1A). The isoforms are classified based on the differential metal composition of the cofactor present in the active site of the nitrogenase, though all three isoforms have a similar protein architecture, consisting of two reductase components and a catalytic component^[3-5]^. Nitrogenases use metalloclusters for electron transfer through the components to the active site cofactor (Fig. 1B). Specifically, upon reduction by a single electron, the reductase component of nitrogenase binds two ATPs to subsequently form a complex with the catalytic component, where one electron is transferred from the [Fe_4_S_4_]-cluster to the P-cluster, an [Fe_8_S_7_]-cluster^[6]^. The electrons flow through the catalytic component from the P-cluster to the active site metallocluster, where the reduction of N_2_ occurs^[7]^. ATP hydrolysis is believed to trigger the release of the reductase component from the catalytic component, with P_*i*_ release being the rate-limiting step ^[8]^. Despite similarities in the mechanisms of N_2_ reduction by the different nitrogenases, the isoforms harbour differential side-activities for reducing non-nitrogen substrates, such as CO_2_ to short-chain hydrocarbons, *i*.*e*. methane, ethene and ethane.^[3, 9-12]^. Notably, the Fe-nitrogenase was shown to have the highest efficiency for reducing CO_2_ compared to the other two nitrogenase isoforms^[13, 14]^.

**Fig. 1.**
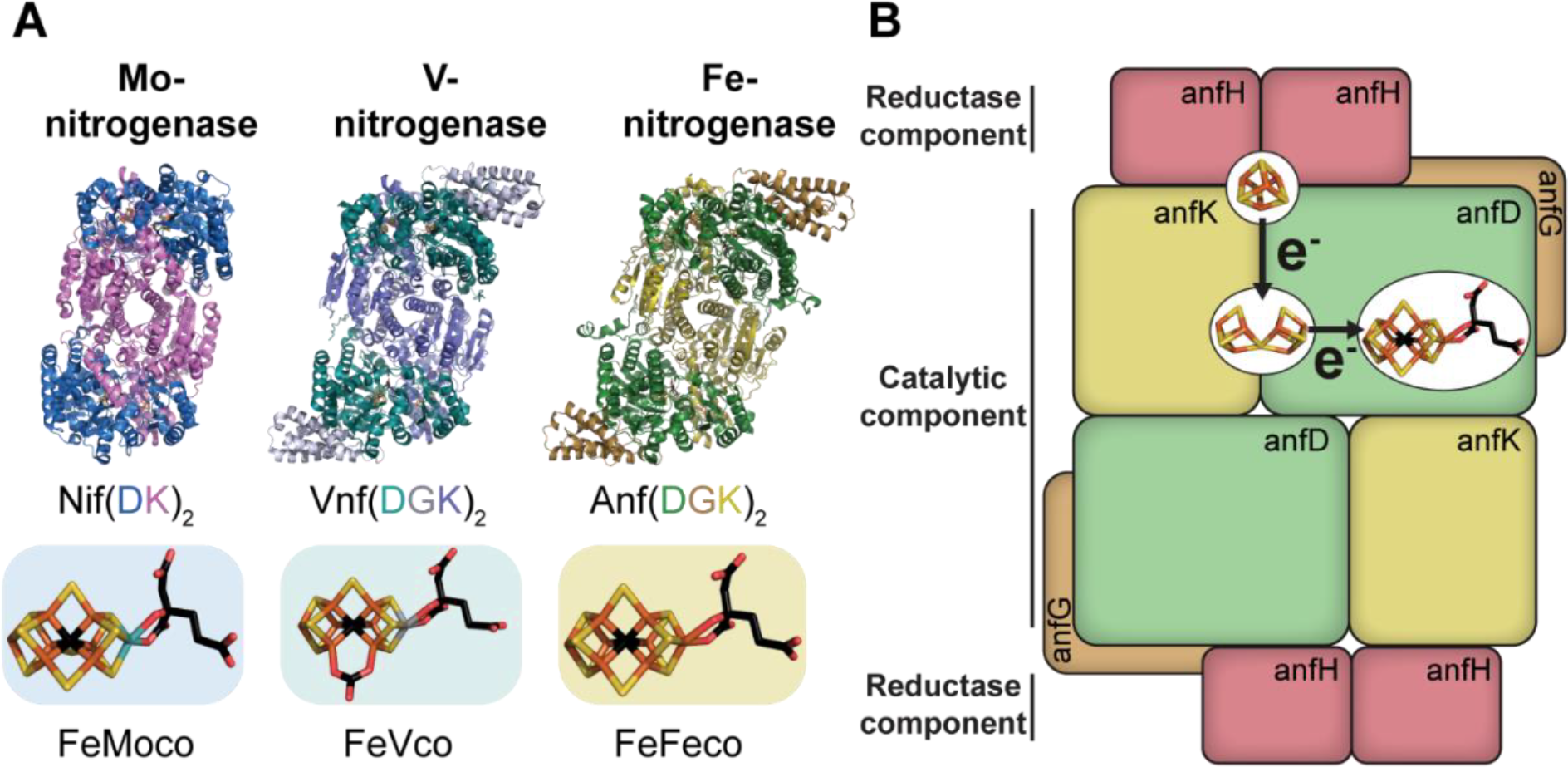
Overview of the three nitrogenase systems, cofactors, electron flow and subunit interactions. **(A)** Structures of the catalytic components and active site cofactors of the three nitrogenase isoforms^[15-18]^. Ribbon presentation of the three nitrogenase catalytic components with subunit annotation below. At the bottom are stick representations of the three active site metalloclusters in coloured boxes. **(B)** Schematic of the electron transfer within the Fe-nitrogenase from the [Fe_4_S_4_]-cluster via the P-cluster to the FeFeco. Black arrows indicate the movement of a single electron. (A-B) The metalloclusters are shown as stick models with carbon in black, iron in orange, oxygen in red, sulphur in yellow, molybdenum in blue and vanadium in grey.

Despite extensive studies examining intramolecular electron transfer within a nitrogenase enzyme, there is relatively limited knowledge about the intermolecular electron transfer to the nitrogenases^[7]^. Only electron transport to Mo-nitrogenases has been investigated previously in *Rhodobacterales*^[19-22]^. The electron transport to either alternative nitrogenase (V- and Fe-nitrogenase) has not been investigated prior^[23]^. A key route of electron transport to Mo-nitrogenase in *R. capsulatus*, which is also expected to be involved in electron transport to the Fe-nitrogenase, is through the Rhodobacter Nitrogen Fixation complex (Rnf complex, Fig. 2). The six structural *rnf* genes are encoded within the N_2_-fixtion gene clusters and are known to be essential for N_2_ fixation by the Mo-nitrogenase^[24, 25]^. Under N_2_-fixing conditions, the Rnf complex is expected to catalyse the reduction of ferredoxins (Fds), utilising energy from a H^+^ or Na^+^ gradient whilst electrons are provided by the oxidation of NADH to NAD^+[26]^. Both, the identity of the electron acceptor for Rnf and the exact redox thermodynamics of Rnf catalysis in *R. capsulatus* remain unknown.

**Fig. 2.**
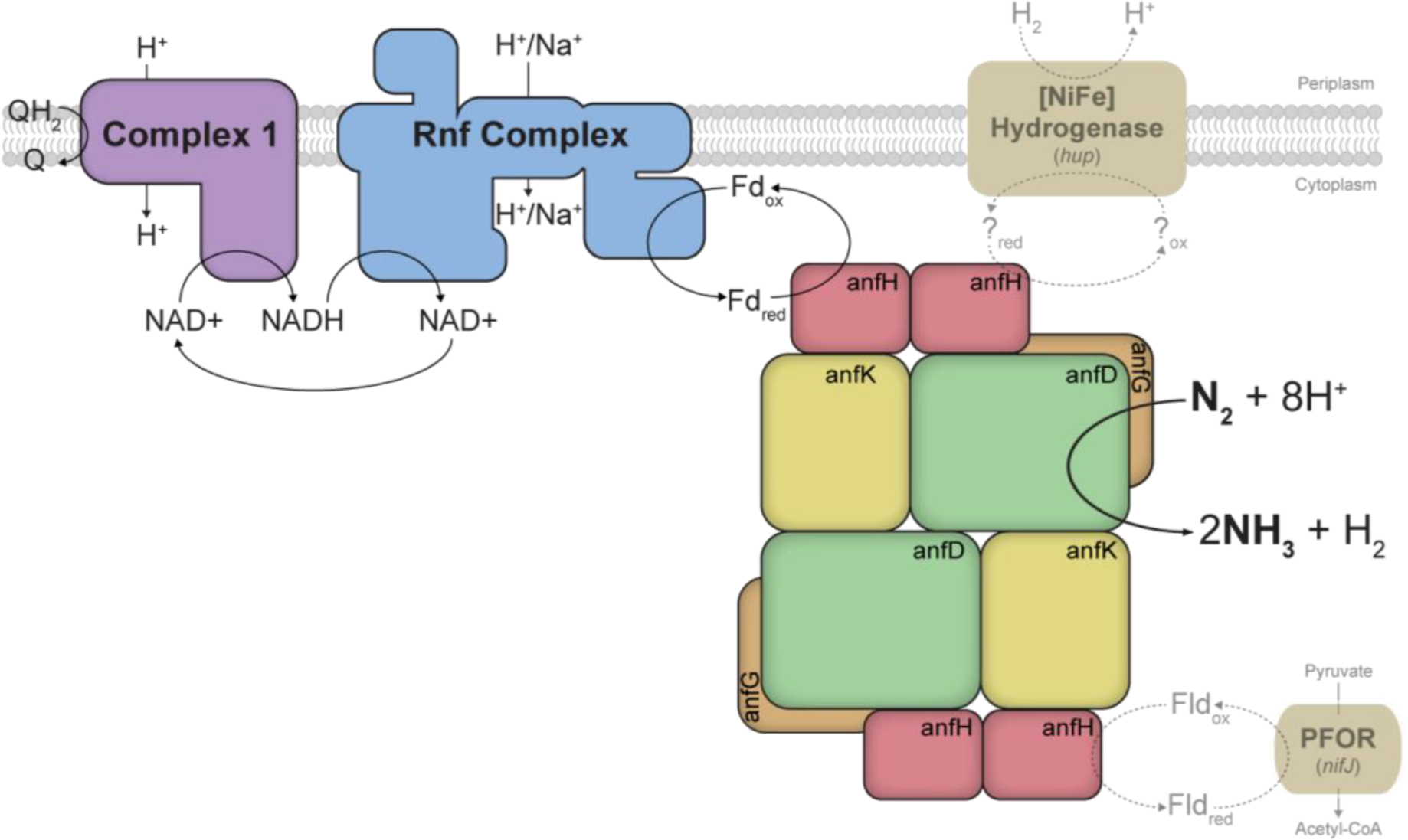
Overview of proposed electron transport routes to the Fe-nitrogenase in *R. capsulatus*. Black arrows indicate the believed prominent route of electrons to the Fe-nitrogenase in *R*. capsulatus. This flow of electrons towards the Fe-nitrogenase is through complex I, the Rnf complex and a Fd of unknown identity. Electrons are believed to flow from the reduced quinone pool in the inner membrane through complex I onto NADH, after which Rnf oxidises NADH to reduce Fd, the reductant of nitrogenase enzymes. Dashed and faint arrows indicate possible alternative electron transfer routes to the Fe-nitrogenase, specifically through the pyruvate:ferredoxin oxidoreductase (PFOR, *nifJ*) and the [NiFe] hydrogenase.

An ample supply of electrons from NADH is necessary for Rnf-mediated Fd reduction in *R. capsulatus*. A key generator of NADH in N_2_-fixing *R. capsulatus* cells is complex I, the NADH:ubiquinone oxidoreductase. Under N_2_-fixing conditions, complex I was shown to mediate reverse electron flow from the membrane quinone pool to reduce NAD^+^ to NADH in the cytosol^[27]^. Deletion of complex I resulted in unviable cells due to an imbalance in the cellular redox homeostasis. Thus, complex I is an essential route for electrons to pass to the Mo-nitrogenase, which is a key electron sink in *R. capsulatus*^[27]^.

A second route of electrons to the Mo-nitrogenase involves the pyruvate-ferredoxin oxidoreductase (NifJ, Fig. 2)^[28]^. NifJ reduces Fds or flavodoxins (Flds) while oxidising pyruvate to acetyl-CoA in anaerobic carbon metabolism and is a critical component of the electron transport pathway to Mo-nitrogenase in other proteobacteria, such as *Klebsiella pneumoniae*^[29]^. *R. capsulatus* NifJ is known to drive the Mo-nitrogenase *in vitro* when combined with purified *R. capsulatus* NifF (flavodoxin 1)^[28]^.

Finally, the uptake [NiFe] hydrogenase HupAB (also called HupSL), which under diazotrophic conditions oxidises H_2_ to H^+^, supports electron transport to nitrogenases (Fig. 2)^[30]^. H_2_ is the main byproduct of nitrogen reduction by nitrogenases and *R. capsulatus* is thought to re-cycle these lost electrons to drive nitrogenases^[31, 32]^. However, the identity of the redox acceptor for the *R. capsulatus* [NiFe] uptake hydrogenase remains unclear. One suggestion is that electrons can move directly from [NiFe] hydrogenase into the ubiquinone pool, where other unknown oxidoreductases can reduce soluble charge carriers to reduce nitrogenases^[33]^.

Soluble charge carriers typically perform the final electron transfer step to nitrogenase enzymes, specifically Fds or Flds^[23]^. Fds contain redox-active Fe-S-clusters, whereas Flds use a flavin mononucleotide (FMN) as the redox cofactor^[26]^. Known charge carriers to nitrogenases are strongly reducing, with very low reduction potentials, thereby allowing the reduction of the [Fe_4_S_4_]-cluster in the subunit interface of the nitrogenase reductase^[34]^. There is considerable interest in the final step of electron delivery to nitrogenases, as it has been suggested to limit N_2_ fixation *in vivo*^[20, 35]^. Interestingly, Fds can have other roles in nitrogen fixation. Some Fds are involved in the maturation of nitrogenase cofactors, donating electrons to assembly proteins such as NifB^[36,37]^.

The genome of *R. capsulatus* contains genes for six Fds (*fdx*), four of which are located on the N_2_ fixation gene clusters, and one Fld (NifF, Fld1), not encoded on the N_2_ fixation gene clusters^[25, 36, 37]^. Only two of the six Fds, FdA and FdE, are essential for *R. capsulatus* growth under non-diazotrophic conditions, shown by the inability to construct viable deletion mutants of the genes *fdxA* and *fdxE*^[38, 39]^. It is unknown which electron carriers are relevant for supplying electrons to the Fe-nitrogenase in *R. capsulatus*, and other diazotrophs, *in vivo*. However, prior work has determined that FdA, FdN and NifF can reduce the *R. capsulatus* Mo-nitrogenase *in vitro* (Table 1).

**Table 1.** Properties of ferredoxins and flavodoxin from *Rhodobacter capsulatus*.

| **Protein** | **Gene** | **Cofactor(s)** | **Donates electrons to Mo-nitrogenase *in vitro*** |
| --- | --- | --- | --- |
| FdA (FdII) | *fdxA\** | [Fe<sub>4</sub>S<sub>4</sub>]- and [Fe<sub>3</sub>S<sub>4</sub>]-cluster | Yes<sup>[40]</sup> |
| FdB (FdIII) | *fdxB* | 2[Fe<sub>4</sub>S<sub>4</sub>]-cluster | No<sup>[41]</sup> |
| FdC (FdIV) | *fdxC* | [Fe<sub>2</sub>S<sub>2</sub>]-cluster | ND |
| FdD (FdV) | *fdxD* | [Fe<sub>2</sub>S<sub>2</sub>]-cluster | No<sup>[42]</sup> |
| FdE (FdVI) | *fdxE\** | [Fe<sub>2</sub>S<sub>2</sub>]-cluster | ND |
| FdN (FdI) | *fdxN* | 2[Fe<sub>4</sub>S<sub>4</sub>]-cluster | Yes<sup>[43]</sup> |
| NifF (FIdI) | *nifF* | FMN | Yes<sup>[44]</sup> |

ND stands for not determined. Other commonly used protein abbreviations shown in brackets. Starred (\*) *fdx* genes could not be deleted from *R. capsulatus* as observed prior<sup>[38, 39]</sup>.

In this study, we characterise the electron transport to the Fe-nitrogenase for the first time. Our experiments show that the individual deletion of *fdxN* or *fdxC* decreases the Fe-nitrogenase-catalysed N_2_ fixation, while the double deletion of Δ*fdxCN* abolishes N_2_ fixation entirely. Next, we started to engineer the electron transport to the Fe-nitrogenase by re-introducing *fdxN* or *fdxC* on plasmids with different copy numbers under the Fe-nitrogenase promoter. *fdxN* complementation was tolerated from both single and high copy number plasmids. In contrast, *fdxC* complementation was only tolerated using a single copy number plasmid, while the high copy number plasmid abolished diazotrophic growth. This information, along with differential proteome shifts for Δ*fdxN* vs Δ*fdxC* strains, showed that FdN and FdC likely have specific roles in *R. capsulatus* and are not redundant in function. Our findings suggest that FdN and FdC have distinct functions in N_2_ fixation, which could originate from the known structural and electrochemical differences^[45]^.

## Results

We set out to identify which soluble charge carriers, *i*.*e*., Fds or Flds in *R. capsulatus*, are associated with diazotrophic growth and N_2_ fixation via the Fe-nitrogenase in a Mo-nitrogenase knockout strain (Δ*nifD, ΔmodABC)*. To achieve this, we used whole-cell proteomics to compare protein abundances between *R. capsulatus* growth on NH_4_^+^ (non-N_2_-fixing conditions), under which N_2_ fixation genes are suppressed^[46]^, and *R. capsulatus* growth with N_2_ as the sole nitrogen source (N_2_-fixing conditions) (Fig. 3).

**Fig. 3.**
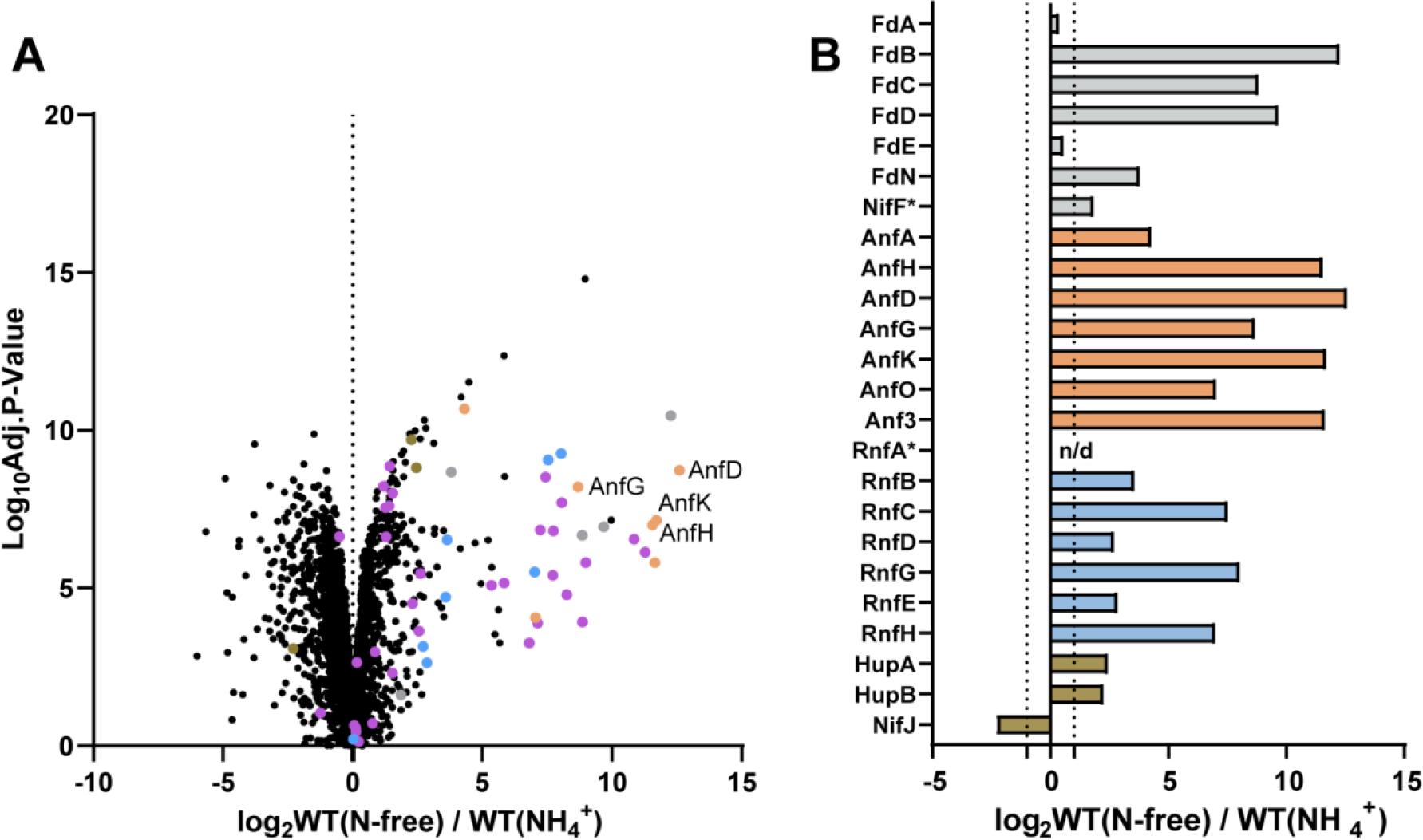
Upregulation of N_2_ fixation proteins in *R. capsulatus* under diazotrophic growth. **(A)** Volcano plot displaying mean intensity log_2_-ratios of N_2_ fixation-related proteins. Fd and Fld proteins (grey), Fe-nitrogenase structural and associated proteins (Anf-protein are labelled in orange), Rnf proteins (light blue), Hup proteins and NifF (brown) and other detected proteins encoded on the N_2_ fixation gene clusters^[25]^ (purple). **(B)** Bar chart displaying average mean intensity log_2_-ratios of highlighted proteins between N_2_-fixing conditions and non-nitrogen fixing conditions. Proteins are colour coded as in (A). Dotted line indicates a log_2_-fold-change of 1 and n/d stands for not detected. (A-B) All proteins with a log_2_-fold-change of >±1 were considered up- or down-regulated between N_2_-fixing conditions (WT (N-free)) and non-nitrogen fixing conditions (WT (NH_4_^+^)). All proteins with a 0.01 P-value (log_10_Adj.P-Value over 2.0) were considered significant, starred proteins (*) did not meet significance criteria. The strain carries an in-frame deletion of Δ*nifD ΔmodABC* to ensure the expression of Fe-nitrogenase genes (*anf)*. Coefficients of variation between 4 independent cultures provided in Table S2.

As expected, most (44) N_2_ fixation-related proteins were upregulated under N_2_-fixing conditions (Fig. 3A), while ten proteins remained similar and only NifJ was downregulated (Table S3). Unsurprisingly, some of the most highly upregulated proteins were Fe-nitrogenase subunits, as diazotrophically grown *R. capsulatus* strictly depends on nitrogenase activity for growth).

Several oxidoreductase proteins implicated in electron transport to nitrogenase were differentially produced under N_2_-fixing conditions (Fig. 3B). Most notably, six of seven Rnf complex subunits were strongly upregulated ^[20, 24]^. The upregulation shows that Rnf production is linked to Fe-nitrogenase-mediated N_2_ fixation and likely has a crucial role in supplying electrons for catalysis. Also, both the small and large subunits of the uptake hydrogenase HupAB had increased expression. HupAB upregulation aligned with prior studies showing increased HupAB activity when nitrogenase is active and producing H_2_, compared to non-N_2_-fixing conditions (NH_4_^+^)^[30]^. Only NifJ was downregulated under N_2_-fixing conditions, supporting prior studies which showed NifJ was produced at similar levels under both N_2_-fixing and non-N_2_-fixing conditions^[28]^. NifJ downregulation implies that it may not be a vital contributor of reduced Fds for Fe-nitrogenase reduction under the tested conditions.

The soluble electron carriers FdC, FdB, FdD and FdN were all significantly upregulated under N_2_-fixing conditions (Fig. 3B). Notably, there was not a significant change in NifF (Fld1) expression under N_2_-fixing conditions despite prior studies showing NifF to drive Mo-nitrogenase *in vitro*^[44]^. The proteome analyses also revealed that expression level of the essential FdA and FdE was almost constant under the tested conditions, suggesting these proteins may not have any additional role under diazotrophic conditions^[38]^.

To evaluate the roles of every upregulated Fd under N_2_-fixing conditions, each Fd was individually and scarlessly deleted from the genome of *R. capsulatus*. Diazotrophic growth was monitored via OD_660_ over time and Fe-nitrogenase activity by the acetylene reduction assay, releasing ethylene and ethane, measured during the exponential growth phase (Fig 4 A, B).

**Fig. 4.**
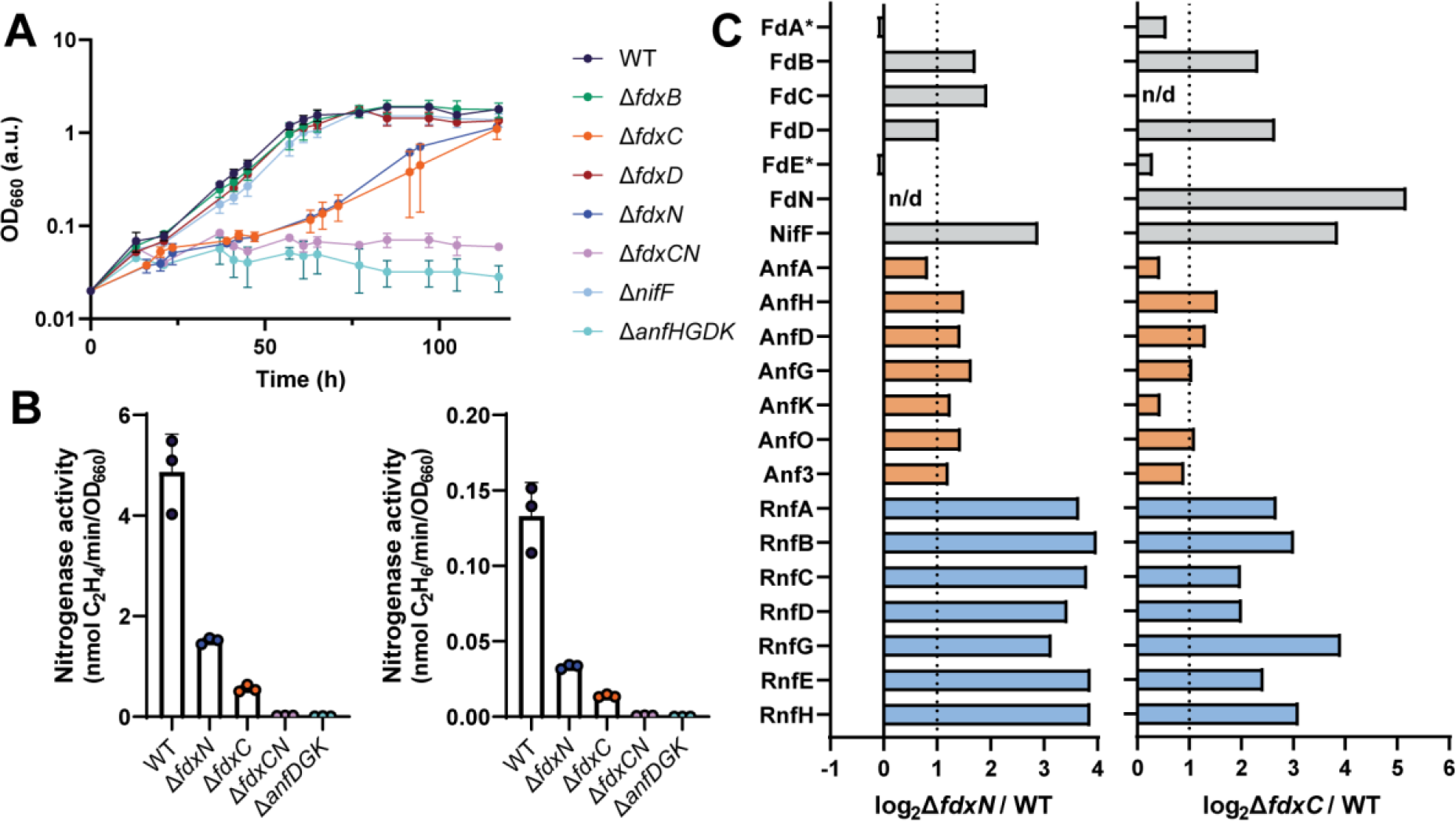
Deletion of *fdxN* and/ or *fdxC* impairs N_2_ fixation by the Fe-nitrogenase in *R. capsulatus*. **(A)** Diazotrophic growth of *R. capsulatus* strains dependent on the Fe-nitrogenase. **(B)** Fe-nitrogenase activity determined via acetylene reduction by *R. capsulatus* strains. All strains, except those with no net OD_660_ increase (Δ*fdxCN, ΔanfHGDK)* were normalised by final OD_660_. **(C)** Bar chart displaying average mean intensity log_2_-ratios of highlighted proteins between Δ*fdxN* and WT (left) and Δ*fdxC* and WT (right). Fd and Fld proteins (grey), Fe-nitrogenase structural and associated proteins (Anf) (orange) and Rnf proteins (light blue). All proteins with a log_2_-fold-change of >±1 were considered up- or downregulated between Δ*fdxN* or Δ*fdxC* strains (Δ*fdxN* or Δ*fdxC*) and the WT. Dotted line indicates a log-fold-change of 1 and n/d stands for not detected. All proteins with a 0.01 P-value (log_10_Adj.P-Value over 2.0) were considered significant, starred proteins (*) did not meet significance criteria. Coefficients of variation between 4 independent cultures provided in Table S4. (A-B) Data are from three independent cultures; error bars represent the standard deviation from the mean. (A-C) All strains carry an in-frame deletion of Δ*nifD ΔmodABC* to ensure the expression of Fe-nitrogenase genes (*anf). R. capsulatus* strains were grown diazotrophically in RCV minimal medium under an N_2_ atmosphere.

Deletion of *fdxN* or *fdxC* slowed diazotrophic growth by 3-4 fold, with doubling times of 28.7 h for Δ*fdxN* and 43.7 h for Δ*fdxC* compared to 10.3 h for the wild-type (Fig. 4A, Table 2). Deletion of both *fdxN* and *fdxC* in the same strain abolished diazotrophic growth by the Fe-nitrogenase, indicating a compounding of the growth issues when both genes are deleted. These results were corroborated by a decrease in Fe-nitrogenase activity of 3-4 fold for the separate deletion strains Δ*fdxN* and Δ*fdxC* and no activity for the double Δ*fdxCN* deletion strain (Fig. 4B). These findings are in agreement with previous Mo-nitrogenase studies, which observed decreased Mo-nitrogenase activity and diazotrophic growth upon deletion of *fdxN* and/or *fdxC*^[22, 39, 47]^.

**Table 2.** Doubling times of *fdx* and *nifF* deletion mutants of *Rhodobacter capsulatus* strains.

| Genotype | Doubling time (h) |
| --- | --- |
| WT | 10.31 ± 0.8 |
| $\Delta fdxB$ | 10.49 ± 1.5 |
| $\Delta fdxC$ | 43.7 ± 21.2 |
| $\Delta fdxD$ | 8.81 ± 0.6 |
| $\Delta fdxN$ | 28.7 ± 4.4 |
| $\Delta fdxCN$ | >100 |
| $\Delta nifF$ | 8.83 ± 0.5 |
| $\Delta anfHGDK$ | >100 |

An exponential Malthusian Growth model was used to determine individual doubling times. Only OD<sub>660</sub> values in the linear range were included. The means and standard deviation of the doubling times were calculated using three biological replicates.

All other deletion strains, Δ*fdxB, ΔfdxD* and Δ*nifF*, had no diazotrophic growth defects. Thus, these genes are not essential for Fe-nitrogenase-mediated N_2_ fixation under photoheterotrophic conditions, or their functions can be complemented by the remaining Fds/ Flds. Our results agree with previous findings for the Mo-nitrogenase, which showed that the interruption of *fdxD, fdxB* and *nifF* did not affect Mo-nitrogenase-based diazotrophic growth^[37, 41, 48]^.

Changes in the proteomes of the individual Δ*fdx* deletion strains were investigated to explore the possible roles of the *fdxN* and *fdxC* gene products, FdN and FdC, respectively. Whole proteome analysis revealed that the proteomes of the Δ*fdxN* and Δ*fdxC* deletion strains changed, relative to the WT under diazotrophic growth, to respond to the loss of one essential *fdx* gene (Fig. S1-2, Table S3-4). There were significant changes in the abundance of soluble charge carriers in both Δ*fdxN* and Δ*fdxC* compared to WT (Fig. 4C). Specifically, deletion of *fdxN* caused an upregulation of FdB, FdD, FdC and NifF. The same trend was observed for *fdxC*, though FdN was the most upregulated soluble charge carrier here, with NifF the second most. *In vitro* studies with the Mo-nitrogenase have shown that both FdN and NifF can donate electrons to the Mo-nitrogenase, though levels of FdN were much higher in cells^[49, 50]^. Perhaps the cells are attempting to push electrons through an alternative electron carrier instead of Fds, when *fdxN or fdxC* are removed.

The observed changes in the abundance of FdN in Δ*fdxC* and NifF in both Δ*fdxC* and Δ*fdxN* may represent a cellular response to overcome the deficits in N_2_ fixation by upregulating the production of proteins capable of reducing the Fe-nitrogenase reductase. Notably, FdN has a log_2_-fold change of over 5 in Δ*fdxC*, revealing a reliance of the cells on FdN when FdC is absent. On the other hand, there was only a log_2_-fold change of 2 for FdC in Δ*fdxN*.

Many further proteins involved in N_2_ fixation were upregulated in both the Δ*fdxN* and Δ*fdxC* strains, the most prominent being Rnf proteins (Fig. 4C). The upregulation of Rnf proteins supported prior work that showed the Rnf complex is the main route of electrons to Mo-nitrogenase in *R. capsulatus*^[20]^. Interestingly, there was no significant change in the abundance of NifJ or HupAB in either Δ*fdxN* or Δ*fdxC* (Fig. S3). It appears that the primary response of the cells is to upregulate the Rnf pathway for electron transport.

Having successfully characterised the *fdx* deletion strains, we aimed to restore WT growth rates by complementing Δ*fdxN* and Δ*fdxC* with plasmids encoding *fdxN* or *fdxC* (Fig. 5, 6). *fdxN* and *fdxC* were separately cloned into a high copy and a single copy plasmid, both under the control of the Fe-nitrogenase promoter (*anfH*), to test which conditions recovered N_2_ fixation.

**Fig. 5.**
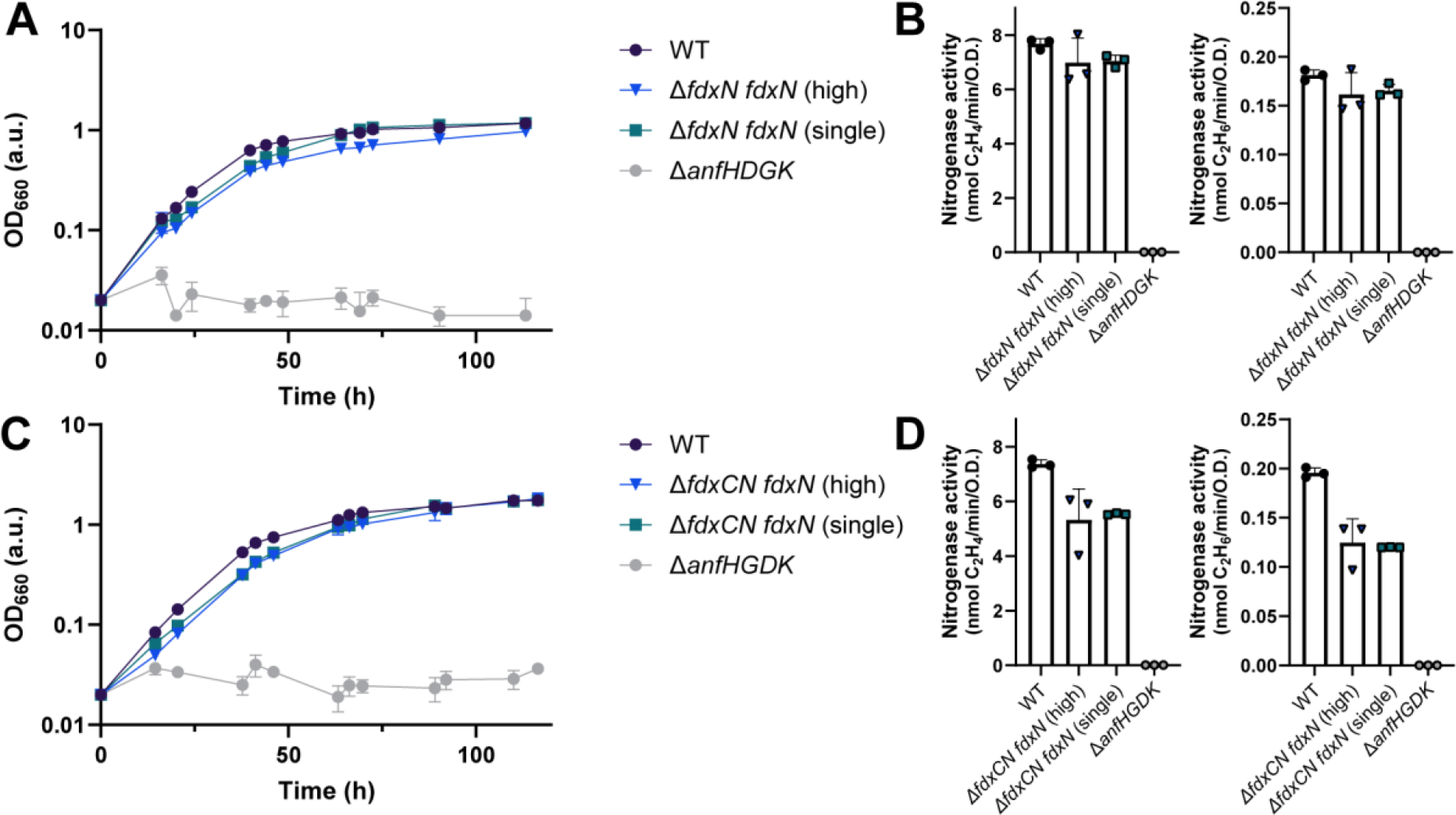
Complementation of *fdxN* recovers N_2_ fixation by the Fe-nitrogenase in *R. capsulatus*. **(A)** Diazotrophic growth of the Δ*fdxN R. capsulatus* complementation strains with *fdxN* on high- and single-copy plasmids. **(B)** Fe-nitrogenase activity determined via acetylene reduction by *R. capsulatus fdxN* complemented Δ*fdxN* strains. **(C)** Diazotrophic growth of the Δ*fdxCN R. capsulatus* complementation strains with *fdxN* on high- and single-copy plasmids. **(D)** Fe-nitrogenase activity determined via acetylene reduction by *R. capsulatus fdxN* complemented Δ*fdxCN* strains. (B, D) All strains except those with no net OD_660_ increase (*ΔanfHGDK)* were normalised by final OD_660_. (A-D) Data are from three independent cultures. Error bars represent the standard deviation from the mean. All strains carry an in-frame deletion of Δ*nifD ΔmodABC* to ensure the expression of Fe-nitrogenase genes (*anf*). *R. capsulatus* strains were grown diazotrophically in RCV minimal medium under an N_2_ atmosphere.

**Fig. 6.**
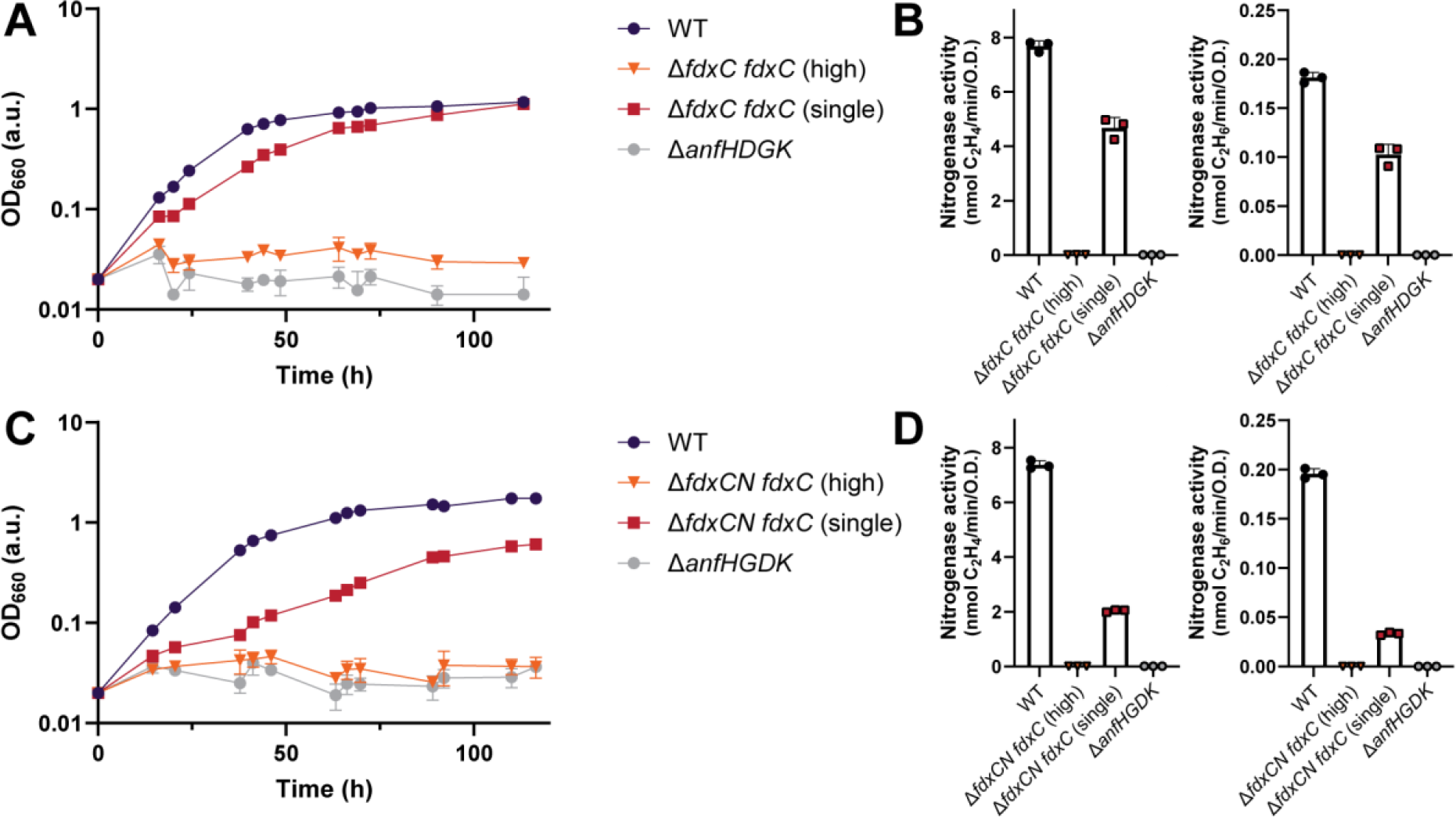
Complementation of *fdxC* on a single copy plasmid recovers N_2_ fixation by the Fe-nitrogenase in *R. capsulatus*. **(A)** Diazotrophic growth of the Δ*fdxC R. capsulatus* complementation strains with *fdxC* on high- and single-copy plasmids. **(B)** Fe-nitrogenase activity determined via acetylene reduction by *R. capsulatus fdxC* complemented Δ*fdxC* strains. **(C)** Diazotrophic growth of the Δ*fdxCN R. capsulatus* complementation strains with *fdxC* on high- and single-copy plasmids. **(D)** Fe-nitrogenase activity determined via acetylene reduction by *R. capsulatus fdxC* complemented Δ*fdxCN* strains. (B, D) All strains except those with no net OD_660_ increase (Δ*fdxC fdxC* (high), *ΔanfHGDK)* were normalised by final OD_660_. (A-D) Data are from three independent cultures. Error bars represent the standard deviation from the mean. All strains carry an in-frame deletion of Δ*nifD ΔmodABC* to ensure the expression of Fe-nitrogenase genes (*anf*). *R. capsulatus* strains were grown diazotrophically in RCV minimal medium under an N_2_ atmosphere.

Expression of *fdxN* under the *anfH* promoter from plasmids recovered diazotrophic growth in both the Δ*fdxN* and the Δ*fdxCN R. capsulatus* strains (Fig. 5A, C). Doubling times of all *fdxN* complementation strains were close to WT, even those in Δ*fdxCN* mutants, at around 10-11 h (Table 3). The doubling time of the Δ*fdxCN fdxN* strain was faster than the Δ*fdxC* strain, which also had one copy of *fdxN* and no copies of *fdxC* but a slower doubling time of around 30 h (Table 2). The faster growth of strain Δ*fdxCN fdxN* could be due to the strength of the *anfH* promoter compared with the *fdxN* native promoter in the genome (the FprA promoter). Differential promoter strengths could cause higher transcription rates of *fdxN* in the Δ*fdxCN fdxN* strain relative to the Δ*fdxC* strain, explaining the doubling time differences. Further, FdN was the most upregulated protein detected in Δ*fdxC*, suggesting it could be important for maintaining Fe-nitrogenase activity in the absence of *fdxC*. Fe-nitrogenase activity was also recovered upon plasmid complementation of *fdxN* in the Δ*fdxN* and Δ*fdxCN R. capsulatus* strains to WT-levels and to around 70% of the WT, respectively (Fig. 5B, D).

**Table 3.** Doubling times of *fdx* complemented *Rhodobacter capsulatus* strains.

| Genotype | Doubling time (h) |
| --- | --- |
| WT | 11.19 ± 0.7 |
| $\Delta fdxN$ *fdxN* (high) | 11.1 ± 0.1 |
| $\Delta fdxN$ *fdxN* (single) | 11.7 ± 0.7 |
| $\Delta fdxC$ *fdxC* (high) | >100 |
| $\Delta fdxC$ *fdxC* (single) | 13.1 ± 1.0 |
| $\Delta fdxCN$ *fdxN* (high) | 10.77 ± 0.2 |
| $\Delta fdxCN$ *fdxN* (single) | 10.91 ± 0.3 |
| $\Delta fdxCN$ *fdxC* (high) | >100 |
| $\Delta fdxCN$ *fdxC* (single) | 23.5 ± 3.0 |
| $\Delta anfHGDK$ | >100 |

An exponential Malthusian Growth model was used to determine individual doubling times. Only OD<sub>660</sub> values in the linear range were included. The means and standard deviation of the doubling times were calculated using three biological replicates.

Interestingly, introducing both multiple copies of *fdxN* (high) and a single copy of *fdxN* (single) recovered growth and nitrogenase activity to similar levels. These results showed that the cells tolerated the expression of *fdxN* under the *anfH* promoter and could be used as a starting point for engineering electron flux to the Fe-nitrogenase. The fact that boosting the number of copies of *fdxN* did not increase nitrogenase activity implies that the bottleneck for electron flow to nitrogenase is at a different stage than the final Fd-mediated electron transfer. Other studies have indicated that the bottleneck is perhaps prior to the reduction of Fd. Specifically, it was shown that upregulation of *rnf* gene expression in *R. capsulatus* resulted in increased Mo-nitrogenase activity^[20]^. Future work could combine the over-expression of the *rnf* genes with the over-expression of *fdxN* to observe if there is increased electron transfer to nitrogenase under these conditions.

Expression of *fdxC* under the *anfH* promoter in Δ*fdxC* and Δ*fdxCN* strains recovered growth and nitrogenase activity but not to WT levels and only when a single copy number plasmid was used (Fig. 6).

Growth and nitrogenase activity were partially recovered when *fdxC* was introduced from a single copy number plasmid into the Δ*fdxC* strain. The doubling time of Δ*fdxC fdxC* (single) was 2 h slower than the WT, increasing from 11 h to 13 h, and nitrogenase activity was only recovered to around 50% of the WT (Fig. 6A, B, Table 3). This observation implies that the genomic context of *fdxC* may be important for functionality. Perhaps the natural *fprA* promoter coordinates the *fdxC* gene expression with other related genes, such as FprA, which might be necessary for the function of FdC.

Growth and nitrogenase activity were only partly recovered when *fdxC* was introduced from a single copy number plasmid into the double Δ*fdxCN* deletion strain (Fig. 6C, D). Doubling times of Δ*fdxCN fdxC* (single) were half as fast as the WT, at 23 h, and nitrogenase activity was around 4-fold less than WT. Both changes in growth and activity were comparable to the Δ*fdxN* deletion strain, which had a similar genotype, being no copies of *fdxN* and one copy of *fdxC*.

Surprisingly, introducing several copies of *fdxC* prevented diazotrophic growth and did not recover nitrogenase activity in both the Δ*fdxC* and Δ*fdxCN* strains. This phenotype was not observed when cultures were grown under ammonium or serine, as the *anfH* promoter was not induced, thus indicating that *fdxC* gene expression was toxic during diazotrophic growth. This toxicity could have been caused by an imbalance in the cellular redox state due to the increased number of FdC proteins per cell. The disruption of stoichiometry to its hypothesised partner protein FprA may have destroyed the cells’ ability to grow diazotrophically^[51]^.

The function of FdC remains undetermined, though prior research provides information into what FdC is likely not doing. Specifically, FdC has a high reduction potential of -275 ± 2mV at pH 7.5 and hence cannot donate electrons to Mo-nitrogenase *in vitro*^[45]^. FdC was suggested to be the physiological electron donor for the N_2_ fixation-related flavoprotein FprA, which was also upregulated in the Δ*fdxC* deletion strain, though the role of FprA remains unknown^[51]^. FdC likely does not act as a direct electron donor to the Fe-nitrogenase but has a different role in the electron transfer pathway. *In vitro* studies with purified FdC and Fe-nitrogenase are necessary to determine the role of FdC.

Overall, our experiments suggest that only FdN and FdC have vital roles in N_2_ fixation by the Fe-nitrogenase in *R. capsulatus*. The results further imply that there is not a high level of functional redundancy between FdN and FdC and their roles are likely distinct.

Building on this information, we wanted to investigate what underpins Fd-mediated electron donation to nitrogenases. Hence, we decided to constructed a phylogeny of all [Fe_4_S_4_]-cluster Fds from *R. capsulatus* (Fig. 7), as FdA and FdN are [Fe_4_S_4_]-cluster Fds known to donate electrons to the Mo-nitrogenase *in vitro* (Table 1)^[40, 43]^. There were three *R. capsulatus* Fds included in the alignment: FdA, FdN and FdB. Despite also being a N_2_ fixation related Fd, the function of FdB is unknown and FdB is incapable of electron transfer to the Mo nitrogenase *in vitro* in spite of its similarity to FdA and FdN^[41]^.The phylogenetic reconstruction provided insight into the genealogical relationships between *R. capsulatus* Fds: FdA, FdB and FdN, and other N_2_ fixation relevant Fds from other model diazotrophs.

**Fig. 7.**
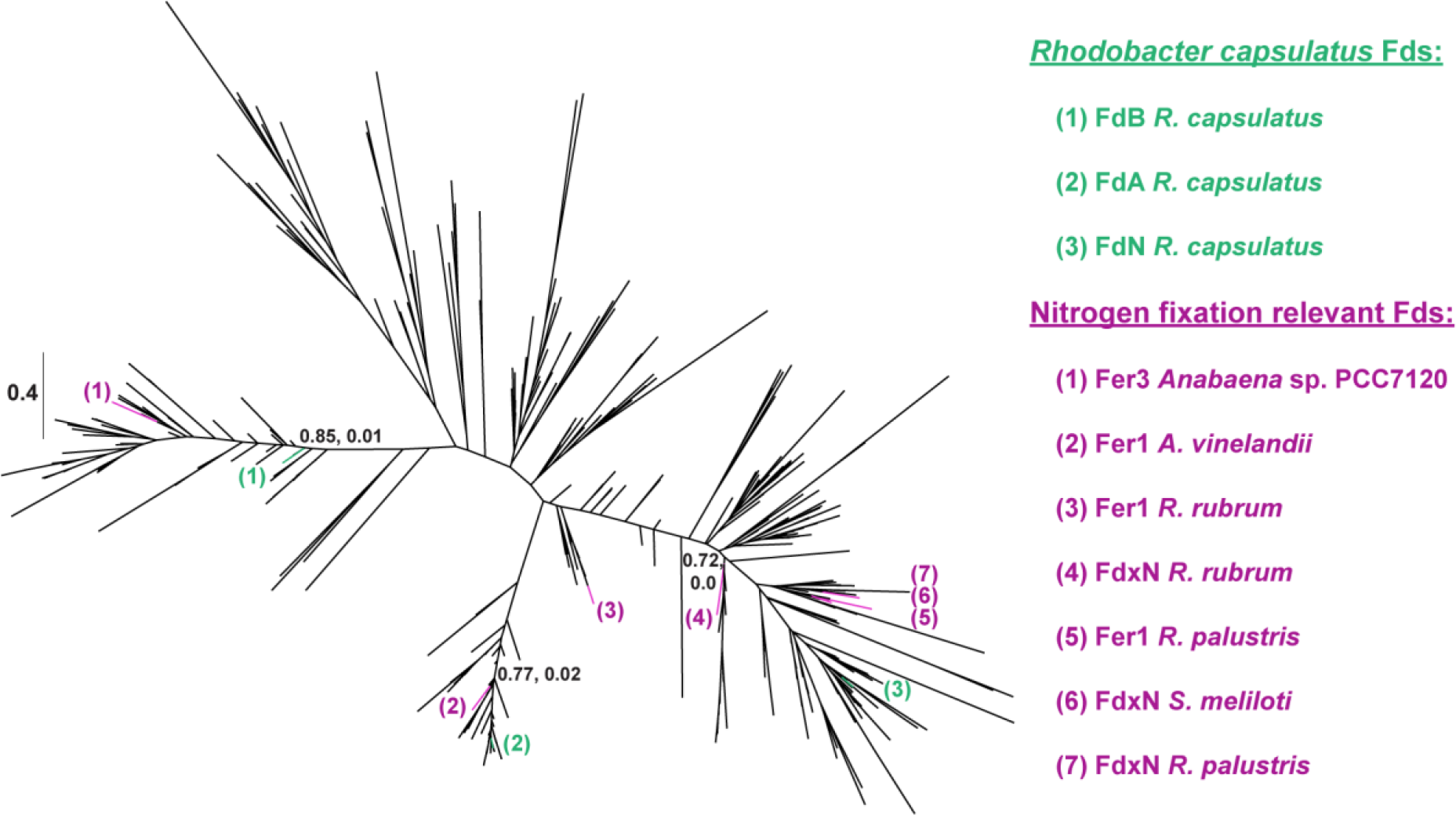
Unrooted phylogenetic tree of [Fe_4_S_4_]-cluster containing ferredoxins from *R. capsulatus*. *R. capsulatus* Fd sequences are coloured in green and numbered as according to the green key on the right. N_2_ fixation-relevant Fd sequences from other model diazotrophs are coloured in purple and numbered according to the purple key on the right. Transfer bootstrap normalised supports (first number) and Felsenstein’s Phylogenetic Bootstrap values (second number) of key internal nodes are indicated on the tree.

As *R. capsulatus* FdN was observed to be important for N_2_ fixation by the Fe-nitrogenase, we were interested if FdN had sequence similarities to other known electron donors to nitrogenases. We observed that FdN branched within other 2[Fe_4_S_4_]-cluster binding proteins, specifically FdN had a high similarity to four other known electron donors to nitrogenase, these being *Rhodopseudomonas palustris* proteins Fer1 and FdN, the *Sinorhizobium meliloti* protein FdxN and the *Rhodospirillum rubrum* protein FdxN^[21, 52, 53]^. Also clustering on the same side of the tree were YhfL-like Fds, which contain 2[Fe_4_S_4_]-clusters and are found in many N_2_-fixing bacteria^[54]^.

On the other hand, Fds grouping closely to FdA were Fds which bind one [Fe_3_S_4_]-cluster and one [Fe_4_S_4_]-cluster. Notably, FdA grouped closley to *Azotobacter vinelandii* protein Fer1, known to drive N_2_ fixation in combination NifF *in vivo*^[55]^. The branching of FdN and FdA near other known electron donors for nitrogenase further supports that both proteins are relevant for N_2_-fixation.

FdB clustered closely to other 2[Fe_4_S_4_]-cluster binding proteins. Interestingly, FdB fell within the same group as a heterocyst Fd from the cyanobacterium *Anabaena* sp. PCC7120 called Fer3, which is involved but not essential to N_2_ fixation^[56]^. The phylogenetic reconstruction did not provide any clear clues as to the function of FdB. However, future studies focusing on sequences at the branching point between FdB, FdN and FdA could give insights into why FdN and FdA can donate electrons to nitrogenases but FdB cannot.

In summary, the phylogenetic reconstruction showed that even among the [Fe_4_S_4_]-cluster binding Fds from *R. capsulatus*, there was significant sequence variation, likely impacting the electrochemical properties of each Fd. Understanding how the biochemistry and biological roles of Fds are impacted by differences in the coordinated metalloclusters and divergent amino acid sequences is critical to understanding these ubiquitous proteins.

## Discussion

Our results are the first investigations into the electron transport to the Fe-nitrogenase. Through whole proteome analysis, the most upregulated proteins under N_2_-fixing conditions by the Fe-nitrogenase were defined and subsequently targetted. Despite deleting the genes for many upregulated soluble charge carriers, only the deletion of *fdxN* or/and *fdxC* disrupted electron transport to the Fe-nitrogenase in *R. capsulatus*. These disruptions were defined by deletion strains growing half as fast diazotrophically and having much lower nitrogenase activity *in vivo*. Proteome analysis of Δ*fdxN* and Δ*fdxC* strains revealed significant upregulation of electron transport proteins, notably NifF, further indicating a disrupted electron transport. Finally, complementation studies of Δ*fdxN* and Δ*fdxC* strains showed distinct patterns for *fdxN* and *fdxC. fdxN* complementation successfully recovered WT phenotypes and was tolerated from both high copy and low copy number plasmids. Whereas *fdxC* complementation only partially recovered WT phenotypes; the Fe-nitrogenase activity was not restored to WT levels and only tolerated a single copy number plasmid. The introduction of several *fdxC* copies was lethal to the *R. capsulatus* cells under N_2_-fixing conditions.

The differences in proteome shifts and complementation patterns between Δ*fdxN* and Δ*fdxC* indicated that FdN and FdC likely have different functions. This conclusion is supported by prior electrochemical characterisations, highlighting key differences in reduction potential, Fe-S-cluster identity and size between FdN and FdC^[41, 50]^. Finally, sequence homology also implied differing functions for FdN and FdC. FdN is more closely related to FdA, the only other Fd shown to donate electrons to the Mo-nitrogenase *in vitro*. In contrast, FdC is closer related to [Fe_2_S_2_]-cluster Fds, *i*.*e*., FdD, which are incapable of reducing nitrogenases^[40]^. Future *in vitro* studies are necessary to characterise electron transfer between Fds and the Fe-nitrogenase. Crucial data, such as comparable reduction potentials and structural data, are required to understand the nature of electron delivery to nitrogenases. Purification of FdN and *in vitro* assays with the Fe-nitrogenase would reveal if FdN can reduce the Fe-nitrogenase reductase, as observed for the Mo-nitrogenase reductase^[43]^. *In vitro* studies would also help elucidate the function of FdC, perhaps either in the electron transfer to the reductase component or in cluster maturation and transfer. Characterising the other proteins within the *fprA* operon may also help reveal the function for FdC, as FdC may be an electron donor or acceptor to one of these proteins^[25]^.

This research has important biotechnological implications. Specifically, characterising the electron transfer pathway to the Fe-nitrogenase is critical for the *in vivo* engineering of *R. capsulatus* for increased gas fixation, either N_2_ or CO_2_. As the bottleneck for N_2_ fixation is believed to be the speed of electron delivery to nitrogenases, electron flux can be altered by targeting Fds, either increasing the number of Fds *in vivo* or engineering the Fds themselves. Additionally, *R. capsulatus* is a biotechnologically relevant host as it generates ATP for N_2_ fixation via anaerobic photophosphorylation using light as an energy source. Due to the availability of light, *R. capsulatus* is a suitable host for the development of sustainable biotechnological processes^[57]^. Finally, our research further defined the minimal gene requirements for N_2_ fixation, which is vital for research looking to transfer a functional nitrogenase systems from diazotrophs into other bacteria or eukaryotes and ultimately plants^[58, 59].^ Future projects will explore the manipulation and optimisation of FdC and FdN to increase electron flux to Fe-nitrogenases, improving N_2_ and CO_2_ conversion.

## Materials and Methods

### Bacterial strains and growth conditions

Chemicals were obtained from Carl Roth (Karlsruhe, Germany) or Tokyo Chemical Industry (Tokyo, Japan), gases from Air Liquide Deutschland GmbH (Düsseldorf, German), enzymes and *E. coli* DH5α from New England Biolabs (Ipswich, United States). Sequencing was performed by Microsynth (Balgach, Switzerland).

*Rhodobacter capsulatus* strains were derivatives of *R. capsulatus* wild-type strain BS85^[48]^. *R. capsulatus* cells were cultured in minimal (RCV) and rich media (PY) according to *Katzke et al*^[60]^.

Diazotrophic growth conditions were photoheterotrophic: RCV minimal media with 30 mM DL-Malic acid^[60]^, 100% N_2_ atmosphere at 0.2 atm at 30°C and under saturated illumination, by LED panels^[9]^ (420nm, 470nm blue lights and 850 nm infrared lights with an output of 12.0 V and light intensity of 80-85 μmol photons m^-2^ s^-1^ per side, or 160-170 total on samples). Non-nitrogen fixing conditions additionally had 10 mM (NH_4_)_2_SO_4_.

*Escherichia coli* (*E. coli*) strains DH5α were cultured in rich media (LB), with 50 μg/mL kamamycin sulfate, at 37°C and 180 rpm.

### *R. capsulatus* deletion strain construction

Deletion strains were constructed using a *sacB* scarless deletion system^[61]^. Flanking homologous regions to each deletion gene were amplified using PCR with Q5 High-fidelity DNA polymerase and cloned into pK18mob-sacB via Golden Gate with BsaI-HF v2. Cloning products were transformed into DH5α and screened via blue-white screening using 200 μg/mL XGAL LB-agar plates. Multiple white colonies were sequenced and a single correct clone was stocked.

Deletion plasmids were transformed into ST18 *E. coli* and subsequently conjugated into *R. capsulatus* as described prior^[60]^. *R. capsulatus* strains were passaged aerobically in PY media for several days before growth anaerobically on 5%-sucrose plates. Counter-selection of single colonies was performed and Kan-sensitive colonies were sequenced. Successful deletion strains were sequenced and stocked.

*E. coli* ST18 cells were grown in the presence of 50 μg/mL kanamycin sulfate and 50 μg/mL 5-aminoleuvelinic acid C_5_H_9_NO_3_.

### *R. capsulatus* complementation strain construction

*fdx* genes were amplified from genomic DNA using PCR with Q5 High-fidelity DNA polymerase. DNA concentrations were determined using a nanodrop (Thermo Scientific, Waltham, United States). PCR fragments were cloned into plasmid via Golden Gate with BsaI-HF v2 or Gibson assembly (New England Biolabs, Gibson Assembly Protocol (E5510)). Cloning products were transformed into DH5α and screened using colony-PCR with DreamTaq polymerase 2X MM. Several promising colonies were sequenced and correct clones were stocked. Expression plasmids were transformed into ST18 *E. coli* and subsequently conjugated into *R. capsulatus* as described prior^[60]^.

The medium/ high copy number plasmid used was pOGG024-Km^R[62]^, with a pBBR1 replication region and kanamycin resistance cassette introduced^[15]^. The single-copy number plasmid was pNMS16, pABC4 replication region and kanamycin resistance cassette (in-press). pOGG024 was a gift from Philip Poole (Addgene plasmid #113991; http://n2t.net/addgene:11991; RRID:Addgene_113991).

### Growth behaviour of *R. capsulatus* deletion and complementation strains

Pre-cultures of *R. capsulatus* deletion strains were inoculated in RCV media with 10mM (NH_4_)_2_SO_4_ and were grown anaerobically for 24 h. Two pre-cultures of *R. capsulatus* complementation strains were inoculated, the first in RCV media with 10 mM serine and grown anaerobically for 24 h and the second with 1 mM serine for grown anaerobically for 24 h. *R. capsulatus* pre-cultures were washed 3-times in N_2_-fixing RCV and OD_660_ was re-measured.

*R. capsulatus* main cultures, inoculated to OD_660_ 0.02, were grown anaerobically in N_2_-fixing RCV media for 5 days. Culture growth was monitored by extracting 100 μL of culture with a nitrogen-flushed needle and OD_660_ was measured using a 96-well plate (Sarstedt AG & Co. KG, Nümbrecht, Germany) and a TECAN Infinite M Nano+ (TECAN, Männedorf, Switzerland). Values were converted to OD_660_ using a calibration factor. Growth of 3 independent cultures were plotted and doubling times were determined using an exponential (Malthusian) growth model in GraphPad prism 9.02 for Windows, GraphPad Software, San Diego, California USA.

*R. capsulatus* deletion strains were grown with 20 μg/ml streptomycin sulfate (Sigma, Darmstadt, Germany) and *R. capsulatus* complementation strains with 50 μg/mL kanamycin sulfate.

### Acetylene reduction activity assay

*In vivo* nitrogenase activity was monitored via acetylene reduction. *R. capsulatus* cultures were prepared as stated before, however after 24 h of diazotrophic growth, 2 ml of each main culture were moved to an empty 20 ml vial in an anaerobic tent (COY Laboratory products, Grass Lake, United States) under argon atmosphere. The headspace of the vial was exchanged with 90% argon, 10% acetylene and incubated under illumination for 4 h at 30°C. Assays were quenched using 100 μL 5 M H_2_SO_4_ (Merck, Rahway, United States). Ethylene and ethane were detected via gas chromatography using the gas chromatograph (GC) PerkinElmer Clarus 690 GC (PerkinElmer, Waltham, United States)^[15]^.

Raw values were converted to nmol via a linear standard curve.

### Proteome analysis

*R. capsulatus* strains were cultured anaerobically until a total OD_660_ of 3 was achieved. Cell samples were prepared by three centrifugation steps and two washing steps with phosphate buffer (3.6 g Na_2_HPO_4_ × 2 H_2_O and 2.6 g KH_2_PO_4_ per litre distilled H_2_O).For protein extraction frozen cell pellets were resuspended in 2% sodium lauroyl sarcosinate (SLS) and heated for 15 min at 90°C. Proteins were reduced with 5 mM Tris(2-carboxyethyl) phosphine (Thermo Fischer Scientific) at 90°C for 15 min and alkylated using 10 mM iodoacetamid (Sigma Aldrich) at 20°C for 30 min in the dark. Proteins were precipitated with a 6-fold excess of ice cold acetone, followed by two methanol washing steps. Dried proteins were reconstituted in 0.2 % SLS and the amount of proteins was determined by bicinchoninic acid protein assay (Thermo Scientific). For tryptic digestion 50 μg protein was incubated in 0.5% SLS and 1 μg of trypsin (Serva) at 30°C over night. Desalted peptides were then analyzed by liquid chromatography-mass spectrometry using an Ultimate 3000 RSLC nano connected to an Exploris 480 mass spectrometer (both Thermo Scientific). Label-free quantification of data independent acquisition raw data was done using DIA-NN. Further details on sample processing, analytical set up and bioinformatics analysis are described in the Supplement.

### Maximum likelihood phylogeny

Protein sequences for *R. capsulatus fdx* genes were collected from Uniprot. Query sequences were as follows: FdA (>WP_013068498.1), FdB (>WP_013068971.1) FdN (>WP_013068980.1). Proteins with sequence similarity to FdA, FdB and FdN were identified by performing individual protein-protein searches in BlastP and tBlastn^[63]^. The experimental databases tool was used to limit sequence redundancy and between 40-60 sequences were selected from each Fd blast. MUSCLE^[64]^ was used to align each set of Fd-related sequences individually, then against each other and this total alignment was viewed in ALiview^[65]^. The alignment was manually refined to remove lineage specific insertions. A maximum likelihood phylogenetic tree was generated using RaxmlHPC-AVX.exe v8.2.10^[66]^ with the PROTGAMMAAUTO, best-scoring model Whelan and Goldman (WAG) matrix, and the Akaike information criterion (aic). The best tree was viewed in FigTree v1.4.4^[67, 68]^. Bootstrap values are TBE normalised supports and Felsenstein’s Phylogenetic Bootstrap values generated from 100 replicates.

## Data availability

Raw data for growth experiments, activity assays, phylogenetic trees and sequence alignments will be deposited on Edmon, the Open Research Data Repository of the Max Planck Society for public access. The mass spectrometry proteomics data has been deposited to the ProteomeXchange Consortium via the PRIDE partner repository with the dataset identifier.

## Acknowledgements

This work was supported by the German Research Foundation (DFG grant 446841743 to J.G.R.). J.G.R. is grateful for generous support from the Max Planck Society. We thank T. Erb and S. Shima for providing equipment essential for carrying out experiments, B. Masepohl and T. Drepper for strains and plasmids and A. Becker for donating the pNMS16 plasmid. We thank N. N Oehlmann, F. V. Schmidt, S. Zhang, S. Pilon and R. Fritsche for the construction of plasmids used in this manuscript. We thank D. Schindler for the Gibson reaction mix. We thank R. Thauer for critical feedback.

## Author contributions

J.G.R. conceived, supervised and acquired funding for this project. H.A. and J.G.R designed and planned experiments. H.A. performed experiments, statistics and bioinformatics. T.G. completed proteome sample preparation, data extraction and provided proteomics expertise. G.H. provided phylogeny expertise. H.A. and J.G.R analysed the data. H.A. and J.G.R. wrote the manuscript, which was reviewed and edited by all authors.

