## Supplemental Information for "Two distinct ferredoxins are essential for nitrogen fixation by the iron nitrogenase in *Rhodobacter capsulatus*"

Running title:
Ferredoxins driving the iron nitrogenase activity

### Supplementary methods:

#### Proteome analysis

*R. capsulatus* strains were cultured anaerobically until a total $OD_{660}$ of 3 was achieved. Cell samples were prepared by three centrifugation steps and two washing steps with phosphate buffer (3.6 g $Na_2HPO_4 \times 2 H_2O$ and 2.6 g $KH_2PO_4$ per litre distilled $H_2O$).For protein extraction frozen cell pellets were resuspended in 2% sodium lauroyl sarcosinate (SLS) and heated for 15 min at 90°C. Proteins were reduced with 5 mM Tris(2-carboxyethyl) phosphine (Thermo Fischer Scientific) at 90°C for 15 min and alkylated using 10 mM iodoacetamid (Sigma Aldrich) at 20°C for 30 min in the dark. Proteins were precipitated with a 6-fold excess of ice cold acetone, followed by two methanol washing steps. Dried proteins were reconstituted in 0.2 % SLS and the amount of proteins was determined by bicinchoninic acid protein assay (Thermo Scientific). For tryptic digestion 50 µg protein was incubated in 0.5% SLS and 1 µg of trypsin (Serva) at 30°C over night.

After digestion, SLS was precipitated by adding a final concentration of 1.5% trifluoroacetic acid (TFA, Thermo Fischer Scientific). Peptides were desalted by using C18 solid phase extraction cartridges (Macherey-Nagel). Cartridges were prepared by adding acetonitrile (ACN), followed by equilibration with 0.1% TFA. Peptides were loaded on equilibrated cartridges, washed with 5% ACN and 0.1% TFA containing buffer and finally eluted with 50% ACN and 0.1% TFA.

Peptides were dried and reconstituted in 0.1% trifluoroacetic acid and then analyzed using liquid-chromatography-mass spectrometry carried out on a Exploris 480 instrument connected to an Ultimate 3000 RSLC nano and a nanospray flex ion source (all Thermo Scientific). Peptide separation was performed on a reverse phase HPLC column (75 µm x 42 cm) packed in-house with C18 resin (2.4 µm; Dr. Maisch). The following separating gradient was used: 94% solvent A (0.15% formic acid) and 6% solvent B (99.85% acetonitrile, 0.15% formic acid) to 25% solvent B over 95 minutes at a flow rate of 300 nl/min, and an additional increase of solvent B to 35% for 25min. MS raw data was acquired in data independent acquisition mode with a method adopted from (https://www.ncbi.nlm.nih.gov/pmc/articles/PMC7124470/). In short, Spray voltage were set to 2.3 kV, funnel RF level at 40, and heated capillary temperature at 275 °C. For DIA experiments full MS resolutions were set to 120.000 at m/z 200 and full MS, AGC (Automatic Gain Control) target was 300% with an IT of 50 ms. Mass range was set to 350–1400. AGC target value for fragment spectra was set at 3000%. 45 windows of 14 Da were used with an overlap of 1 Da. Resolution was set to 15,000 and IT to 22 ms. Stepped HCD collision energy of 25, 27.5, 30 % was used. MS1 data was acquired in profile, MS2 DIA data in centroid mode.

Analysis of DIA data was performed using DIA-NN version 1.8 (https://www.ncbi.nlm.nih.gov/pmc/articles/PMC6949130/) using a uniprot protein database from *Rhodobacter capsulatus*. Full tryptic digest was allowed with two missed cleavage sites, and oxidized methionines and carbamidomethylated cysteins. Match between runs and remove likely interferences were enabled. The neural network classifier was set to the single-pass mode, and protein inference was based on genes. Quantification strategy was set to any LC (high accuracy). Cross-run normalization was set to RT-dependent. Library generation was set to smart profiling. DIA-NN outputs were further evaluated using the SafeQuant (https://pubmed.ncbi.nlm.nih.gov/23017020/ & https://analyticalsciencejournals.onlinelibrary.wiley.com/doi/10.1002/pmic.201300135) script modified to process DIA-NN outputs.

**<u>Supplementary materials:</u>**

**Table S1: Strains, plasmids and primers used in this study.**

**Strains:**

| Strain | Genotype | Reference |
|---|---|---|
| ***R. capsulatus* strains** | | |
| BS85 Δ*modABC* | Wild-type: Δ*nifD*::Sp mutant (Δ*nifDK*) Δ*modABC* | [1] |
| Δ*fdxB* | Δ*nifD*::Sp mutant (Δ*nifDK*) Δ*modABC* Δ*fdxB* | This study |
| Δ*fdxC* | Δ*nifD*::Sp mutant (Δ*nifDK*) Δ*modABC* Δ*fdxC* | This study |
| Δ*fdxD* | Δ*nifD*::Sp mutant (Δ*nifDK*) Δ*modABC* Δ*fdxD* | This study |
| Δ*fdxN* | Δ*nifD*::Sp mutant (Δ*nifDK*) Δ*modABC* Δ*fdxN* | This study |
| Δ*nifF* | Δ*nifD*::Sp mutant (Δ*nifDK*) Δ*modABC* Δ*nifF* | This study |
| Δ*fdxCN* | Δ*nifD*::Sp mutant (Δ*nifDK*) Δ*modABC* Δ*fdxCN* | This study |
| Δ*anfHGDK* | Δ*nifD*::Sp mutant (Δ*nifDK*) Δ*modABC* Δ*anfDGK*::*GmR* | This study |
| ***E. coli* strains** | | |
| DH5α | Φ80*lac*ZΔM15 Δ(*lac*ZYA-*arg*F) U169 *rec*A1 *end*A1 *hsd*R17($r_K^-$, $m_K^+$) *pho*A *sup*E44 *thi*-1 *gyr*A96 *rel*A1 $\lambda^-$ | Thermo Fisher Scientific Inc. (Waltham, USA) catalogue #18265017 |
| ST18 | RP4-2 *Tc*::*Mu Km*::*Tn7* Δ*hemA* mutant | [2] |

**Plasmids:**

| Plasmid | Relevant Features | Reference |
|---|---|---|
| pK18mob-SacB | Suicide plasmid, *sacB*, *oriT* (mobilisable), *lacZα* cassette, $Km^R$ | [3] |
| pK18mob-SacB2 | Suicide plasmid, *sacB*, *oriT* (mobilisable), *lacZα* cassette for golden gate cloning (BsaI), $Km^R$ | This study |
| pK18mob-SacB2-Δ*fdxB* | Suicide plasmid, *sacB*, *oriT* (mobilisable), in-frame Δ*fdxB*, $Km^R$ | This study |
| pK18mob-SacB2-Δ*fdxC* | Suicide plasmid, *sacB*, *oriT* (mobilisable), in-frame Δ*fdxC* into BsaI site of MCS, $Km^R$ | This study |
| pK18mob-SacB2-Δ*fdxD* | Suicide plasmid, *sacB*, *oriT* (mobilisable), in-frame Δ*fdxD*, $Km^R$ | This study |
| pK18mob-SacB2-Δ*fdxN* | Suicide plasmid, *sacB*, *oriT* (mobilisable), in-frame Δ*fdxN* into BsaI site of MCS, $Km^R$ | This study |
| pK18mob-SacB2-Δ*nifF* | Suicide plasmid, *sacB*, *oriT* (mobilisable), in-frame Δ*nifF* into BsaI site of MCS, $Km^R$ | This study |
| pK18mob-SacB2-Δ*fdxCN* | Suicide plasmid, *sacB*, *oriT* (mobilisable), in-frame Δ*fdxCN*, $Km^R$ | This study |
| pOGG024-$Km^R$ | Broad-host range plasmid, pBBR1 with *oriT* (mobilisable), *lacZα* cassette for golden gate cloning (BsaI), $Km^R$ | [4] |
| pOGG024-2-$Km^R$ | Broad-host range plasmid, pBBR1 with *oriT* (mobilisable), *lacZα* cassette for golden gate cloning (BsaI), AnfH promoter, $Km^R$ | This study |
| pOGG024-2-$Km^R$ *fdxN* | Broad-host range plasmid, pBBR1 with *oriT* (mobilisable), *fdxN* cloned into BsaI site, AnfH promoter, $Km^R$ | This study |
| pOGG024-2-$Km^R$ *fdxC* | Broad-host range plasmid, pBBR1 with *oriT* (mobilisable), *fdxC* cloned into BsaI site, AnfH promoter, $Km^R$ | This study |
| pNMS16 | Broad-host range plasmid, repABC4 with *oriT* (mobilisable), $Km^R$ | (in-press) |
| pNMS16-2 | Broad-host range plasmid, repABC4 with *oriT* (mobilisable), *lacZα* cassette for golden gate cloning (BsaI), $Km^R$ | This study |

| | | |
|---|---|---|
| pNMS16-3 | Broad-host range plasmid, repABC4 with *oriT* (mobilisable), AnfH promoter, $Km^R$ | This study |
| pNMS16-3-*fdxN* | Broad-host range plasmid, repABC4 with *oriT* (mobilisable), *fdxN* cloned into MCS, AnfH promoter, $Km^R$ | This study |
| pNMS16-3-*fdxC* | Broad-host range plasmid, repABC4 with *oriT* (mobilisable), *fdxC* cloned into MCS, AnfH promoter, $Km^R$ | This study |

**Primers:**

| Primer | Sequence (5'-3') | Purpose |
|---|---|---|
| LacZ-F | TTACACTCTTCCGTAGGGGGTTACTCTAGGG | pOGG024 LacZ and MCS for Golden Gate into pNMS16 via Gibson |
| LacZ-R | GTCAAGTGGGATGCGTTCGGTCAAGGTTCTG | pOGG024 LacZ and MCS for Golden Gate into pNMS16 via Gibson |
| pK18mob-SacB-F | GACCGAACGCATCCCACTTGACCGAGATACA | Gibson cloning LacZ and MCS for Golden Gate into pNMS16 |
| pK18mob-SacB-R | ACCCCCTACGGAAGAGTGTAAGCGGGTTGA | Gibson cloning LacZ and MCS for Golden Gate into pNMS16 |
| anfHDGK-F | CACCACAGGTCTCGGGAGGCAGCCCGTTCGGAATTCC | GoldenGate anfH promoter into pNMS16-3 |
| anfHGDK-R | CACCACAGGTCTCGAGCGTCACCACACGTTGAGGATCC | GoldenGate anfH promoter into pNMS16-3 |
| anfHp-F | CAGCCCGTTCGGAATTCC | Gibson cloning anfH promoter into pOGG024-$Km^R$ |
| anfHp-R | CATATGTATATCTCCTTCTTAAAGTTAAACAAAAT | Gibson cloning anfH promoter into pOGG024-$Km^R$ |
| pOGG024-$Km^R$-F | CTTTAAGAAGGAGATATACATATGTGAGACCGCAGCTGGCACG | Gibson cloning anfH promoter into pOGG024-$Km^R$ |
| pOGG024-$Km^R$-R | GGAATTCCGAACGGGCTGCAAATAAAACGAAAGGCTCAGTCG | Gibson cloning anfH promoter into pOGG024-$Km^R$ |
| Δ*fdxB*-UF | GATTCAGGTCTCCCGGTCTGCTCGTCGATCTTCATCAGG | Amplify *fdxB* upstream homologous region from BS85 |
| Δ*fdxB*-UR | GATCTAGGTCTCTTACGTCGGGCTTCTGGTCGAAAAG | Amplify *fdxB* upstream homologous region from BS85 |
| Δ*fdxB*-DF | GATTCAGGTCTCCTTACATCAATCGGGGGCAAAGCAG | Amplify *fdxB* downstream homologous region from BS85 |
| Δ*fdxB*-DR | GATTCAGGTCTCCACCGGGGCCGAGAATTGT | Amplify *fdxB* downstream homologous region from BS85 |
| pK18mob-SacB-F | CGTAATAGCGAAGAGGCCCG | Amplification of pK18mob-SacB for Gibson cloning |
| pK18mob-SacB-R | GTAAAACGACGGCCAGTGC | Amplification of pK18mob-SacB for Gibson cloning |
| pK18mob-SacB-Qu-F | ACAATTCTCCTGCTCGTCGAT | Frame-shift correction via Quickchange in pK18mob-SacB-Δ*fdxB* |

| | | |
|---|---|---|
| pK18mob-SacB-Qu-R | GGAGAATTGTATCGGCTGCG | Frame-shift correction via Quickchange in pK18mob-SacB-Δ*fdxB* |
| Δ*fdxC*-UF | GATCTAGGTCTCTATGGTTATCTGACCCATACGCGGC | Amplify *fdxC* upstream homologous region from BS85 |
| Δ*fdxC*-UR | GATCTAGGTCTCCCTCGATGATGCGGGTTCC | Amplify *fdxC* upstream homologous region from BS85 |
| Δ*fdxC*-DF | GATCTAGGTCTCTCGAGGAAATGCGGGTGCTGGAGG | Amplify *fdxC* downstream homologous region from BS85 |
| Δ*fdxC*-DR | GATCTAGGTCTCCGGAGCCAGAAGATTGTTGCCCGAC | Amplify *fdxC* downstream homologous region from BS85 |
| Δ*fdxD*-UF | GATCTAGGTCTCCAGGCCACCGCGTAGATGGTCTTGTC | Amplify *fdxD* upstream homologous region from BS85 |
| Δ*fdxD*-UR | GATCTAGGTCTCCTACGGGTTGATCAGCGTCGACAGC | Amplify *fdxD* upstream homologous region from BS85 |
| Δ*fdxD*-DF | GATCTAGGTCTCCTTACAGTCCACATCGTCATAGGCG | Amplify *fdxD* downstream homologous region from BS85 |
| Δ*fdxD*-DR | GATCTAGGTCTCCGCCTGCCAGTTCATCCCG | Amplify *fdxD* downstream homologous region from BS85 |
| Δ*fdxN*-UF | GATCTAGGTCTCTCGCACGCATTGGTCGGGCAGAC | Amplify *fdxN* upstream homologous region from BS85 |
| Δ*fdxN*-UR | GATCTAGGTCTCTATGGCAGGGGCTGAAGATGCGTC | Amplify *fdxN* upstream homologous region from BS85 |
| Δ*fdxN*-DF | GATCTAGGTCTCCGGAGATGCACGTCGAGCACCTG | Amplify *fdxN* downstream homologous region from BS85 |
| Δ*fdxN*-DR | GATCTAGGTCTCCTGCGAAGGCGAGCATGAC | Amplify *fdxN* downstream homologous region from BS85 |
| Δ*fdxCN*-UF | GATCTAGGTCTCCCTCGATGATGCGGGTTCC | Amplify *fdxCN* upstream homologous region from BS85 |
| Δ*fdxCN*-UR | GATCTAGGTCTCTTACGGATCACCCCCTTCCTGGATC | Amplify *fdxCN* upstream homologous region from BS85 |
| Δ*fdxCN*-DF | GATCTAGGTCTCCTTACATGCACGTCGAGCACCTG | Amplify *fdxCN* downstream homologous region from BS85 |
| Δ*fdxCN*-DR | GATCTAGGTCTCCCGAGTGCGAAGGCGAGCATGAC | Amplify *fdxCN* downstream homologous region from BS85 |
| pK18mob-SacB-F | GATTTAGGTCTCTGTAAAACGACGGCCAGTGC | GoldenGate cloning of *fdxD* or *fdxCN* homolgous regions into pK18mob-SacB |

| | | |
|---|---|---|
| pK18mob-SacB-R | GATTTAGGTCTCTCGTAATAGCGAAGAGGCCCG | GoldenGate cloning of *fdxD* or *fdxCN* homolgous regions into pK18mob-SacB |
| Δ*nifF*-UF | GATCTAGGTCTCTATGGATATCAAAGGCCCCGGCG | Amplify *nifF* upstream homologous region from BS85 |
| Δ*nifF*-UR | GATCTAGGTCTCTTGATCCTCATCGTCGAACATGTCC | Amplify *nifF* upstream homologous region from BS85 |
| Δ*nifF*-DF | GATCTAGGTCTCTATCAGGACAATCAGGCGGC | Amplify *nifF* downstream homologous region from BS85 |
| Δ*nifF*-DR | GATCTAGGTCTCTGGAGTGAAGGAATCCGACTGGTGC | Amplify *nifF* downstream homologous region from BS85 |
| *fdxN*-Gi-F | CTTTAAGAAGGAGATATACATATGGCCATGAAGATCGATCCCGA | Amplify *fdxN* for Gibson cloning into pNMS16-3 |
| *fdxN*-Gi-R | CAACAGGAGTCCAAGAGCGTTACGCCGCCGGGTTG | Amplify *fdxN* for Gibson cloning into pNMS16-3 |
| *fdxC*-Gi-F | TTTACTTTAAGAAGGAGATATACATATGGACAAGGCCACACTGACGTT | Amplify *fdxC* for Gibson cloning into pNMS16-3 |
| *fdxC*-Gi-R | TCAACAGGAGTCCAAGAGCGTCAGGCGGGGCGAACC | Amplify *fdxC* for Gibson cloning into pNMS16-3 |
| pNMS16-3-F | CGCTCTTGGACTCCTGTTGA | Gibson cloning of *fdxN* or *fdxC* into pNMS16-3 |
| pNMS16-3-R | CATATGTATATCTCCTTCTTAAAGTTAAACAAAAT | Gibson cloning of *fdxN* or *fdxC* into pNMS16-3 |
| *fdxN*-Go-F | GATCTAGGTCTCTAGCGTTACGCCGCCGGGTTGATG | Amplify *fdxN* for GoldenGate into pOGG024-2-$Km^R$ |
| *fdxN*-Go-R | GATCTAGGTCTCCTATGGCCATGAAGATCGATCCCGA | Amplify *fdxN* for GoldenGate into pOGG024-2-$Km^R$ |
| *fdxC*-Go-F | GATCTAGGTCTCTAGCGTCAGGCGGGGCGAACCTT | Amplify *fdxC* for GoldenGate into pOGG024-2-$Km^R$ |
| *fdxC*-Go-R | GATCTAGGTCTCCTATGATGGACAAGGCCACACTGACG | Amplify *fdxC* for GoldenGate into pOGG024-2-$Km^R$ |

**Table S2: Whole proteome analysis of *R. capsulatus* grown under non-$N_2$-fixing conditions ($NH_4^+$) and $N_2$-fixing conditions (N-free).** (Excel file attached).

**Table S3: $Log_2$ ratios of $N_2$ fixation proteins between *R. capsulatus* grown under $N_2$-fixing conditions (N-free) and non-$N_2$-fixing conditions ($NH_4^+$).**

| **Protein code** | **Gene name** | **N-free.log$_2$ratio** | **-log$_{10}$_pValue.N-free** |
|---|---|---|---|
| D5ANJ7 | *anfD* | 12.59872 | 8.727144 |
| D5ARX7 | *fdxB* | 12.27127 | 10.466 |
| D5ANJ9 | *anfK* | 11.70715 | 7.152174 |
| D5ANK2 | *RCAP_rcc00591* | 11.65023 | 5.812046 |

| | | | |
|---|---|---|---|
| D5ANJ6 | *anfH* | 11.56207 | 6.997151 |
| D5ANI3 | *nifH* | 11.27484 | 6.136328 |
| D5ARX9 | *RCAP_rcc03277* | 10.85964 | 6.545178 |
| D5ANI4 | *fdxD* | 9.679919 | 6.939873 |
| D5ARW4 | *RCAP_rcc03262* | 8.987266 | 5.808905 |
| D5ARY0 | *nifX* | 8.859228 | 3.927786 |
| D5ARY7 | *fdxC* | 8.845235 | 6.675845 |
| D5ANJ8 | *anfG* | 8.689296 | 8.207057 |
| D5ARY1 | *nifN* | 8.256918 | 4.784761 |
| D5ARW5 | *RCAP_rcc03263* | 8.068804 | 7.712417 |
| D5ARZ3 | *rnfG* | 8.044382 | 9.263996 |
| D5ARX3 | *nifU2* | 7.744715 | 6.812609 |
| D5ARX8 | *RCAP_rcc03276* | 7.726194 | 5.40691 |
| D5ARZ1 | *rnfC* | 7.536894 | 9.055173 |
| D5ARY8 | *fprA* | 7.432053 | 8.5205 |
| D5ARX4 | *RCAP_rcc03272* | 7.228431 | 6.836491 |
| D5ARY2 | *nifE* | 7.128616 | 3.889748 |
| D5ANK0 | *anfO* | 7.047755 | 4.065219 |
| D5ARZ5 | *rnfH* | 7.017223 | 5.510083 |
| D5ARX2 | *nifS* | 6.814573 | 3.258705 |
| D5ANH9 | *rpoN* | 5.842829 | 5.168062 |
| D5ARX0 | *nifW* | 5.350851 | 5.090256 |
| D5ANJ5 | *anfA* | 4.307349 | 10.68301 |
| D5ARY6 | *fdxN* | 3.79751 | 8.671147 |
| D5ARY4 | *rnfF* | 3.636934 | 6.530411 |
| D5ARZ0 | *rnfB* | 3.584759 | 4.715786 |
| D5ARZ4 | *rnfE* | 2.869379 | 2.6354 |
| D5ARZ2 | *rnfD* | 2.713729 | 3.150655 |
| D5ANH7 | *nifB1* | 2.613498 | 5.460614 |
| D5ANI2 | *nifD* | 2.559534 | 3.638374 |
| D5APG0 | *hupA* | 2.451492 | 8.811083 |
| D5ARY3 | *RCAP_rcc03281* | 2.308116 | 4.51212 |
| D5APG1 | *hupB* | 2.262473 | 9.702227 |
| P52967 | *nifF* | 1.856026 | 1.610574 |
| P09431 | *ntrB* | 1.541781 | 8.008296 |
| D5ANI6 | *cowN* | 1.529626 | 2.30679 |
| P09432 | *ntrC* | 1.427897 | 8.858428 |
| D5AKV9 | *modA2* | 1.408857 | 7.605173 |
| D5ANH8 | *nifA1* | 1.288746 | 6.626321 |
| D5ANH2 | *mopA* | 1.263358 | 7.549659 |
| D5ANH6 | *modD* | 1.185609 | 8.222177 |

| | | | |
|---|---|---|---|
| D5AKV8 | *modB2* | 0.857103 | 2.98255 |
| D5ARX1 | *nifV* | 0.767107 | 0.711768 |
| D5ANI1 | *nifK* | 0.235134 | 0.137075 |
| D5AUA6 | *ntrX* | 0.165516 | 2.642961 |
| D5AKW0 | *modC2* | 0.127946 | 0.454067 |
| D5AUA5 | *ntrY* | 0.11615 | 0.570263 |
| D5ARW3 | *RCAP_rcc03261* | 0.057427 | 0.651148 |
| D5ARY9 | *rnfA* | 0.022949 | 0.207997 |
| D5ANH1 | *mopB* | -0.51132 | 6.632272 |
| D5ANI0 | *nifU1* | -1.23967 | 1.048143 |
| D5AU34 | *nifJ* | -2.28244 | 3.084097 |

All proteins with a log-fold-change of >±1 were considered up- or down-regulated between $N_2$-fixing conditions (WT (N-free)) and non-$N_2$ fixing conditions (WT ($NH_4^+$)). All proteins with a 0.01 P-value ($log_{10}$Adj.P-Value over 2.0) were considered significant. Boxes coloured in green are upregulated proteins, boxes in orange are downregulated proteins and boxes not coloured represent proteins that either are not up- or down- regulated or do not meet the 0.01 P-value cut-off. n/a stands for not applicable.

**Table S4: Whole proteome analysis of WT, Δ*fdxN* and Δ*fdxC* *R. capsulatus* strains grown under $N_2$-fixing conditions.** (Excel file attached).

**Table S5: Log$_2$ ratios of $N_2$ fixation proteins between *R. capsulatus* Δ*fdxN* or Δ*fdxC* strains and the *R. capsulatus* WT.**

| **Protein code** | **Gene name** | **Δ*fdxN*. log$_2$ratio** | **Δ*fdxC*. log$_2$ratio** | **-log$_{10}$_pValue. Δ*fdxN*** | **-log$_{10}$_pValue. Δ*fdxC*** |
|---|---|---|---|---|---|
| D5ARZ0 | *rnfB* | 3.998439 | 3.027834 | 9.19467 | 9.239084 |
| D5ARZ4 | *rnfE* | 3.885121 | 2.440782 | 7.046392 | 6.272526 |
| D5ARZ5 | *rnfH* | 3.87851 | 3.116919 | 8.86022 | 9.117162 |
| D5ARZ1 | *rnfC* | 3.81805 | 2.007708 | 7.632947 | 6.38165 |
| D5ARY9 | *rnfA* | 3.668276 | 2.692474 | 5.858928 | 6.280726 |
| D5ARZ2 | *rnfD* | 3.451044 | 2.02617 | 6.755926 | 5.853414 |
| D5ARZ3 | *rnfG* | 3.156771 | 3.933726 | 11.36347 | 11.94346 |
| P52967 | *nifF* | 2.90767 | 3.871691 | 3.493402 | 4.201193 |
| D5ARW4 | *RCAP_rcc03262* | 2.102174 | 2.491553 | 8.052484 | 8.983435 |
| D5ANI2 | *nifD* | 2.049606 | 2.697424 | 6.784045 | 8.173014 |
| D5ARY7 | *fdxC* | 1.949947 | n/a | 7.054317 | 9.579414 |
| D5ARX4 | *RCAP_rcc03272* | 1.839633 | 2.043956 | 6.623682 | 8.086195 |
| D5ARY4 | *rnfF* | 1.827542 | 2.151687 | 6.008059 | 7.963533 |
| D5ARX9 | *RCAP_rcc03277* | 1.810358 | 2.6456 | 6.698211 | 8.144624 |

| | | | | | |
|---|---|---|---|---|---|
| D5ARX8 | *RCAP_rcc03276* | 1.781268 | 2.718973 | 7.896731 | 9.972142 |
| D5ARX7 | *fdxB* | 1.730894 | 2.338911 | 7.390378 | 5.618482 |
| D5ARY8 | *fprA* | 1.698857 | 1.390543 | 9.248192 | 8.597107 |
| D5ANH7 | *nifB1* | 1.667965 | 2.165186 | 5.687121 | 6.819651 |
| D5ANJ8 | *anfG* | 1.658196 | 1.327339 | 4.931956 | 5.164582 |
| D5ARW5 | *RCAP_rcc03263* | 1.561616 | 2.14193 | 6.018241 | 6.937882 |
| D5ARX0 | *nifW* | 1.544861 | 1.776224 | 5.661919 | 6.866204 |
| D5ANJ6 | *anfH* | 1.51508 | 1.553795 | 5.930304 | 6.645611 |
| D5ANI3 | *nifH* | 1.499192 | 2.024093 | 5.430483 | 6.650974 |
| D5ANH9 | *rpoN* | 1.461493 | 1.81334 | 7.179225 | 7.045798 |
| D5ANK0 | *anfO* | 1.456562 | 1.117837 | 4.504814 | 5.400112 |
| D5ANJ7 | *anfD* | 1.451124 | 1.070575 | 5.183646 | 5.506334 |
| D5ARX3 | *nifU2* | 1.334243 | 2.048841 | 6.343032 | 8.711973 |
| D5ANI5 | *RCAP_rcc00574* | 1.26887 | -1.97086 | 2.555368 | 1.494688 |
| D5ANJ9 | *anfK* | 1.265425 | 0.462979 | 6.1094 | 4.527944 |
| D5ANK2 | *RCAP_rcc00591* | 1.228654 | 0.917518 | 5.132341 | 5.127066 |
| D5ARY2 | *nifE* | 1.181117 | 1.426355 | 5.994891 | 6.197274 |
| D5ARY0 | *nifX* | 1.165747 | 1.830177 | 5.883956 | 8.474927 |
| D5ARX2 | *nifS* | 1.128982 | 1.373777 | 4.826705 | 5.090009 |
| D5ARY1 | *nifN* | 1.092217 | 1.479451 | 7.018562 | 8.523274 |
| D5ANI4 | *fdxD* | 1.045993 | 2.66351 | 5.026649 | 10.22016 |
| D5AKV8 | *modB2* | 0.949781 | -0.13845 | 2.719426 | 0.608573 |
| D5AU34 | *nifJ* | 0.932884 | 0.563581 | 1.56201 | 0.062862 |
| D5ARX1 | *nifV* | 0.900374 | 0.892005 | 1.421565 | 1.07132 |
| D5ANJ5 | *anfA* | 0.845423 | 0.454307 | 5.526384 | 3.873301 |
| D5ARY3 | *RCAP_rcc03281* | 0.766416 | -0.21199 | 1.145381 | 0.291941 |
| D5AKW0 | *modC2* | 0.725204 | -0.67275 | 3.229507 | 4.379903 |
| D5ARW3 | *RCAP_rcc03261* | 0.622173 | 1.069767 | 4.260913 | 7.233544 |
| D5ANI6 | *cowN* | 0.604531 | -2.81857 | 0.87265 | 4.595866 |
| D5ANI1 | *nifK* | 0.472799 | 1.226359 | 0.550185 | 1.28607 |
| D5ANI0 | *nifU1* | 0.387758 | 3.356679 | 0.548121 | 3.347977 |
| D5AKV9 | *modA2* | 0.359114 | 0.167948 | 2.200597 | 1.790392 |
| D5AUA5 | *ntrY* | 0.170983 | -0.17455 | 0.712654 | 0.738389 |
| D5AUA6 | *ntrX* | -0.01533 | -0.02527 | 0.126457 | 0.241044 |
| D5ANH2 | *mopA* | -0.14889 | -0.55089 | 1.457109 | 5.424188 |
| D5ASC6 | *mod* | -0.15287 | -0.17444 | 2.307833 | 2.492609 |
| D5ANH8 | *nifA1* | -0.16688 | -1.12742 | 1.013266 | 6.500072 |
| D5ANH1 | *mopB* | -0.19284 | -0.88574 | 2.346441 | 6.935616 |

| | | | | | |
|---|---|---|---|---|---|
| P09431 | *ntrB* | -0.20457 | 0.399407 | 3.251092 | 3.810276 |
| P09432 | *ntrC* | -0.22451 | -0.30686 | 3.136265 | 3.8755 |
| D5ANH6 | *modD* | -0.27382 | -0.19504 | 3.625535 | 2.871633 |
| D5APG0 | *hupA* | -0.71449 | -0.51455 | 5.927237 | 5.191983 |
| D5APG1 | *hupB* | -0.72956 | -0.61754 | 6.664283 | 6.304395 |
| D5ARY6 | *fdxN* | n/a | 5.193566 | 2.473119 | 2.53946 |

All proteins with a log-fold-change of >±1 were considered up- or downregulated between Δ*fdxN* or Δ*fdxC* strains (Δ*fdxN* or Δ*fdxC*) and the WT (WT). All proteins with a 0.01 P-value ($log_{10}$Adj.P-Value over 2.0) were considered significant. All strains were grown diazotrophically in RCV minimal medium under an $N_2$ atmosphere ($N_2$-fixing conditions). Boxes coloured in green are upregulated proteins, boxes in orange are downregulated proteins and boxes not coloured represent proteins that either are not up- or down- regulated or do not meet the 0.01 P-value cut-off. n/a stands for not applicable.

**Table S6. Sequences used for phylogenetic reconstruction of ferredoxins from *Rhodobacter capsulatus*.**

| Protein | Sequence (FASTA) |
|---|---|
| FdA | >WP_013068498.1 ferredoxin family protein [Rhodobacter capsulatus] MTYVVTDNCIACKYTDCVEVCPVDCFYEGENTLVIHPDECIDCGVCE PECPADAIRPDTEPGMEDWVEFNRTYASQWPVITIKKDPMPDHKKY DGETGKREKYFSPNPGTGD |
| FdB | >WP_013068971.1 ferredoxin III, nif-specific [Rhodobacter capsulatus] MMPTVAYTRGGAEYTPVYLMKIDEQKCIGCGRCFKVCGRDVMSLHG LTEDGQVVAPGTDEWDEVEDEIVKKVMALTGAENCIGCGACARVCP SECQTHAALS |
| FdN | >WP_013068980.1 ferredoxin FdxN [Rhodobacter capsulatus] MAMKIDPELCTSCGDCEPVCPTNAIAPKKGVYVINADTCTECEGEH DLPQCVNACMTDNCINPAA |

Sequences were retrieved from the UniProt database[5].

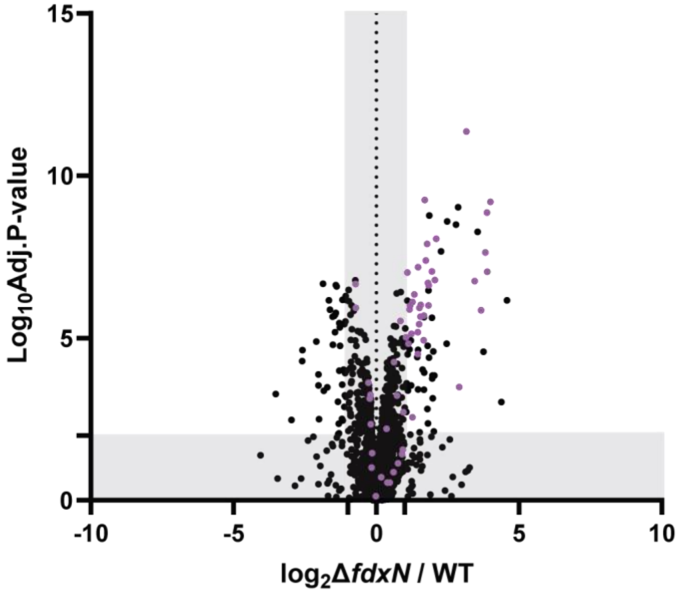


**Fig S1. Up-regulation of nitrogen fixation-related proteins in *R. capsulatus* ∆*fdxN* strain relative to *R. capsulatus* WT.** Volcano plot displaying abundance ratios of proteins between *R. capsulatus* ∆*fdxN* strains and *R. capsulatus* WT. $N_2$ fixation-related proteins are shown in purple. All proteins with a $log_2$-fold-change of >±1 were considered up- or down-regulated between *R. capsulatus* ∆*fdxN* and *R. capsulatus* WT strains. All proteins with a 0.01 P-value ($log_{10}$Adj.P-Value over 2.0) were considered significant. Grey boxes highlight regions of $log_2$fold values of -1> n <1 and $log_{10}$P-values of <2. Coefficients of variation between 4 independent cultures provided in Table S4. All strains carry an in-frame deletion of ∆*nifD* ∆*modABC* to ensure the expression of Fe-nitrogenase genes (*anf*). *R. capsulatus* strains were grown diazotrophically in RCV minimal medium under an $N_2$ atmosphere.

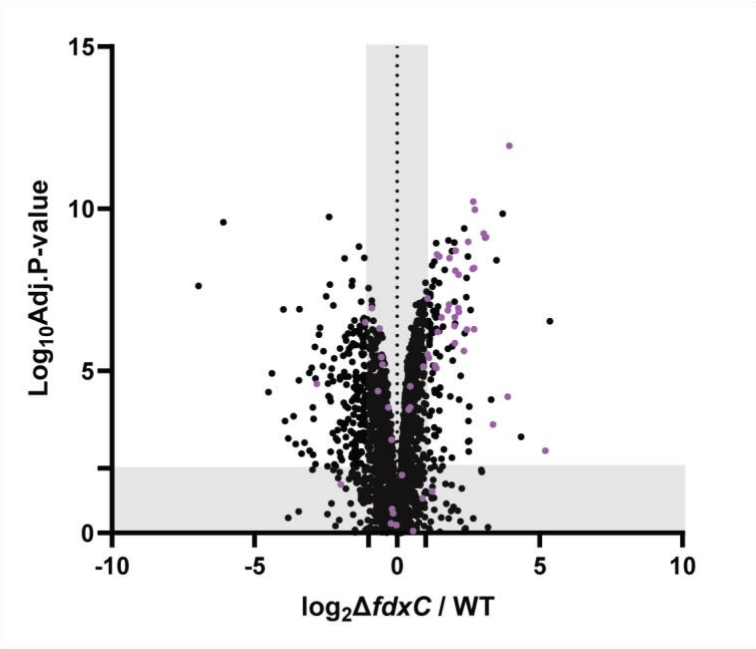


**Fig. S2. Up-regulation of nitrogen fixation-related proteins in *R. capsulatus* ∆*fdxC* strain relative to *R. capsulatus* WT.** Volcano plot displaying abundance ratios of proteins between *R. capsulatus* ∆*fdxC* strains and *R. capsulatus* WT. $N_2$ fixation-related proteins are shown in purple. All proteins with a $log_2$-fold-change of >±1 were considered up- or down-regulated between *R. capsulatus* ∆*fdxC* and *R. capsulatus* WT strains. All proteins with a 0.01 P-value ($log_{10}$Adj.P-Value over 2.0) were considered significant. Grey boxes highlight regions of $log_2$fold values of -1> n <1 and $log_{10}$P-values of <2. Coefficients of variation between 4 independent cultures provided in Table S4. All strains carry an in-frame deletion of ∆*nifD* ∆*modABC* to ensure the expression of Fe-nitrogenase genes (*anf*). *R. capsulatus* strains were grown diazotrophically in RCV minimal medium under an $N_2$ atmosphere.

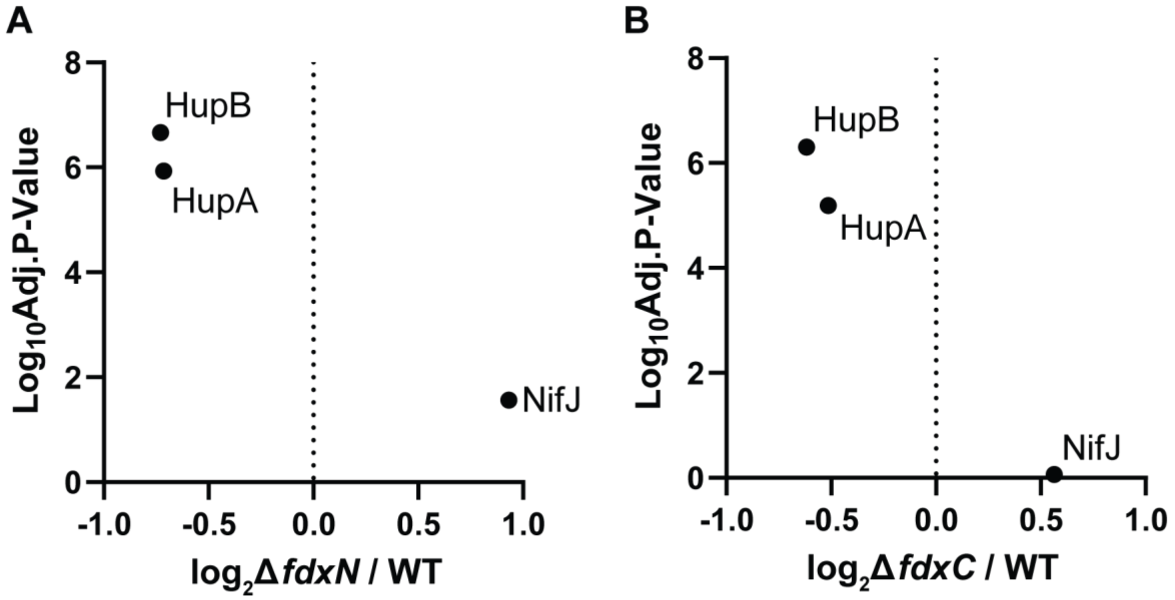


**Fig. S3. No significant abundance changes in NifJ, HupA and HupB in ∆*fdxN* and ∆*fdxC* vs WT. (A)** Scatter plot displaying abundance ratios of NifJ, HupA and HupB in ∆*fdxN* compared to the WT. **(B)** Scatter plot displaying abundance ratios of NifJ, HupA and HupB in ∆*fdxC* to compared to the WT. (A-B) All strains carry an in-frame deletion of ∆*nifD* ∆*modABC* to ensure the expression of Fe-nitrogenase genes (*Anf*). *R. capsulatus* strains were grown diazotrophically in RCV minimal medium under an $N_2$ atmosphere. All proteins with a 0.01 P-value ($log_{10}$Adj.P-Value over 2.0) were considered significant. All proteins with a $log_2$-fold-change of value of >±1 were considered over or under produced, between ∆*fdxN* or ∆*fdxC* to WT. Coefficients of variation between 4 independent cultures provided in Table S4.
